# Extensions of BLUP models for genomic prediction in heterogeneous populations: Application in a diverse switchgrass sample

**DOI:** 10.1101/124081

**Authors:** Guillaume P. Ramstein, Michael D. Casler

## Abstract

Genomic prediction is a useful tool to accelerate genetic gain in selection using DNA marker information. However, this technology usually relies on models that are not designed to accommodate population heterogeneity, which results from differences in marker effects across genetic backgrounds. Previous studies have proposed to cope with population heterogeneity using diverse approaches: (i) either ignoring it, therefore relying on the robustness of standard approaches; (ii) reducing it, by selecting homogenous subsets of individuals in the sample; or (iii) modelling it by using interactive models. In this study we assessed all three possible approaches, applying existing and novel procedures for each of them. All procedures developed are based on deterministic optimizations, can account for heteroscedasticity, and are applicable in contexts of admixed populations. In a case study on a diverse switchgrass sample, we compared the procedures to a control where predictions rely on homogeneous subsamples. Ignoring heterogeneity was often not detrimental, and sometimes beneficial, to prediction accuracy, compared to the control. Reducing heterogeneity did not result in further increases in accuracy. However, in scenarios of limited subsample sizes, a novel procedure, which accounted for redundancy within subsamples, outperformed the existing procedure, which only considered relationships to selection candidates. Modelling heterogeneity resulted in substantial increases in accuracy, in the cases where accounting for population heterogeneity yielded a highly significant improvement in fit. Our study exemplifies advantages and limits of the various approaches that are promising in various contexts of population heterogeneity, e.g. prediction based on historical datasets or dynamic breeding.

## INTRODUCTION

Genomic prediction has proved a useful tool to predict genetic merit in plant and animal breeding (Hayes *et al.* 2009a, Lorenz *et al.* 2011). This technology consists of learning relationships between DNA markers and phenotypes, which arise from the non-random association (linkage disequilibrium; LD) between DNA markers and causal genetic variants having direct effects on the trait studied (Meuwissen *et al.* 2001). Typical genomic prediction models, including genomic BLUP (GBLUP; VanRaden 2008, Hayes *et al.* 2009b) or Bayesian linear regression (BLR) models (Meuwissen *et al.* 2001, Gianola *et al.* 2009), assume that the effects of causal variants are linear and purely additive, so estimated effects do not capture any dependence on context, arising for example from interactions of causal variants with environmental or genetic backgrounds. Initially, genomic prediction models have been proposed for applications in populations that are relatively homogeneous with respect to LD patterns and interactions involving causal variants (Meuwissen *et al.* 2001). In such situations, increasing the size of the calibration set (CS) – the set of individuals used to estimate the model’s parameters – would typically benefit accuracy of the models (Lorenzana & Bernardo 2009, VanRaden *et al.* 2009). However, in practice, increasing the CS size may often involves calibrating prediction models on individuals with inconsistent LD patterns and/or backgrounds, which may result in reduced accuracy (Wientjes *et al.* 2016). This issue will arise in the typical situation where an initially homogeneous CS is augmented with individuals from extraneous populations, that is, multi-population – or (in the animal literature) multi-breed – calibration (Lund *et al.* 2014). Recently, studies in both plant and animal breeding have assessed the usefulness of combining populations from different genetic backgrounds in genomic prediction. In general, one or two of the following approaches were studied: (i) single-population prediction in a multi-population context (ignoring population heterogeneity); (ii) instance selection (reducing population heterogeneity); and (iii) multi-population prediction (modelling population heterogeneity).

In single-population prediction (*SPM*), the simulation study of De Roos *et al.* (2009) suggested that adding an extraneous population to a CS may benefit prediction accuracy if the added population is not too dissimilar (in terms of divergence time) from the initial CS. These authors also suggested that high enough marker density could prevent prediction accuracy from decreasing, even in cases of strong divergence between populations. Consistently, most empirical studies of multi-population calibration with high marker density, based on single-population BLR and/or GBLUP, have reported little or no gain in accuracy under strong population structure (Lehermeier *et al.* 2015, Jarquín *et al.* 2016, Hayes *et al.* 2009c, Erbe *et al.* 2012). In contrast, only a few studies have reported substantial increases in accuracy from multi-population calibration in similar conditions (Technow *et al*. 2013, Daetwyler *et al.* 2012). Interestingly, Habier *et al.* (2013) suggested that increasing CS size may reduce prediction accuracy in GBLUP, even in a single-population context, due to accumulated noise in the larger genomic relationship matrix, especially when many relationship coefficients are small. In an attempt to increase the accuracy of genomic relationship estimation, Endelman and Jannink (2012) and Müller *et al.* (2015) have proposed regularization methods, which proved especially useful when marker density was low. However, the regularization methods proposed in these two studies did not account for potential population structure in the genomic relationship matrix, which would naturally arise in a multi-population context.

In instance selection (*IS*) – or training set design/optimization – only a subset of the available individuals is selected to make up the CS. Studies have generally focused on a scenario of limited phenotyping resources, where the sample of individuals was searched for an optimal CS of pre-determined size. The CS searches in these studies were either stochastic or deterministic. Stochastic searches in this context have consisted in randomly choosing a CS to maximize some measure of prediction accuracy for the selection candidates, using either random exchange algorithms (Rincent *et al.* 2012, Isidro *et al.* 2014, Rutkoski *et al.* 2015) or genetic algorithms (Akdemir *et al.* 2015), which were compared to purely random sampling as a baseline. Studies of this type have used selection criteria such as the prediction error variance (Henderson 1984) or the mean coefficient of determination (Laloë 1993) as a measure of accuracy, and have generally concluded that stochastic searches guided by one of these criteria performed better than random sampling. One disadvantage of stochastic searches is that they are computationally intensive, so deterministic searches may be preferred in some scenarios (e.g., when sample size is large). This second type of searches has typically involved choosing the set of individuals so as to maximize some measure of relatedness between the CS and the selection candidates (Clark *et al.* 2012, Lorenz and Smith 2015). The contribution of such relatedness to accuracy has been asserted by simulation studies (Pszczola *et al.* 2012, Wientjes *et al.* 2013). However, Pszczola *et al.* (2012) also suggested that accuracy was negatively impacted by relationships within the CS, for a given CS size (probably owing to redundancy in information). To our knowledge, no deterministic search in genomic prediction has accounted for that trade-off involving relationships.

In multi-population prediction (*MPM*), studies have proposed to fit, to the whole set of available individuals, models that were capable of accommodating population heterogeneity explicitly. This type of models includes multi-trait GBLUP models, with “traits” corresponding to population backgrounds (Karoui *et al.* 2012, Carillier *et al.* 2014, Lehermeier *et al.* 2015), and random regression models based on markers interacting with discrete population cluster coefficients (de los Campos *et al.* 2015, with a BLR model). To our knowledge, the implementation of these methods has not been adapted to contexts of admixture, where population structure variables are continuous. Furthermore, when calibration involves many populations, the increase in model complexity of these methods will make them computationally intractable and statistically inefficient. Parsimonious multi-population models, based on only a few parameters to capture population heterogeneity, have also been proposed (Zhou *et al.* 2014, Heslot and Jannink 2015). In presence of many populations, such models are more practical and potentially more useful than multi-trait and random interaction models. Also, since they generally assume some underlying basis for population heterogeneity (e.g., inconsistency in LD patterns), they may generate insight about the causes of marker-by-population interactions.

In this study, we investigated the usefulness of *SPM, IS* and *MPM* for coping with population heterogeneity. We present a general framework for the application of existing and novel methods under each of these three approaches. All these procedures were compared to a control procedure (*Target*) where the CS includes only the individuals from the same population as the selection candidates, as is typically done to avoid dealing with population heterogeneity. We applied the procedures to the analysis of three traits (plant height, heading date, and standability) in switchgrass (*Panicum virgatum* L.), an herbaceous biomass crop showing good promise for bioenergy production (Sanderson *et al.* 1996, Perlack *et al.* 2005, Perlack *et al.* 2011, Langholtz *et al.* 2016). This species is characterized by an extensive diversity which make it particularly suitable for studying population heterogeneity (Casler 2012). The sample under study comprised seven population clusters from two diverse panels, assayed in the Midwestern region of the United States, which represent differentiation by ecotype (upland or lowland), geographical origin (latitudinal and longitudinal gradients) and ploidy level (tetraploid or octoploid). This sample exemplified the heterogeneity of data available for practical applications of genomic prediction, which pose both opportunities (by increased sample sizes) and challenges (by inconsistencies in marker effects across populations) for breeding based on DNA markers.

In this study, we did not fit BLR models (usually based on Markov chain Monte Carlo optimizations), since we focused on deterministic methods for model fit and considered only models based on best linear unbiased predictors (BLUP). Besides, we did not model potential genotype-by-environment interactions in the sample and did not accommodate potential heteroscedasticity across populations (with respect to errors or marker effects). Finally, we did not study any context of across-population prediction, defined as prediction based on a CS that completely excludes the population of selection candidates. Further research would be needed to address the aforementioned issues.

The present work describes promising methods for increasing accuracy and robustness of predictions in situations where heterogeneous data sources are combined, for example when the CS incorporates data from historical trials (Dawson *et al.* 2013, Rutkoski *et al.* 2015) or from multiple generations of a dynamic breeding program (Sallam *et al.* 2015, Auinger *et al.* 2016).

## MATERIAL AND METHODS

Table 1 describes the abbreviations about panels, traits and procedures used to this study.

**Table 1.** Abbreviations specific to this study

| Type | Abbreviation | Description |
| --- | --- | --- |
| Panels | BP | Breeding panel |
|  | AP | Association panel |
| Traits | PH | Plant height |
|  | HD | Heading date |
|  | St | Standability |
| Sets of individuals in validation | CS | Calibration set |
|  | TS | Testing set |
| Optimization approaches | *SPM* | Single-population model |
|  | *IS* | Instance selection |
|  | *MPM* | Mixed-population model |
| Optimization procedures | *Target* | Control procedure |
|  | *SPM-GRM* | *SPM*: genomic relationship matrix (Reference) |
|  | *SPM-GLASSO* | *SPM*: graphical LASSO (Proposed) |
|  | *IS-Rel* | *IS*: relationships (Reference) |
|  | *IS-QP* | *IS*: quadratic programming (Proposed) |
|  | *MPM-Mixture* | *MPM*: mixture model (Reference) |
|  | *MPM-Matérn* | *MPM*: Matérn kernel model (Proposed) |

Reference: pre-existing procedure, under a given optimization approach. Proposed: procedure proposed in this study, under a given optimization approach.

### Panels and populations

In this study, two multi-population panels were assayed and considered together in one sample. The first panel was the breeding panel (BP) described in Ramstein *et al.* (2016), comprising two tetraploid breeding populations of half-sib families: WS4U-C2, which consisted of 137 half-sib families derived from a diverse upland-ecotype pool of 162 plants (Casler *et al.* 2006), and Liberty-C2, which consisted of 110 half-sib families derived from the lowland-upland cultivar Liberty (Casler and Vogel 2014). The second panel was the association panel (AP) described in Lu *et al.* (2013) and Evans *et al.* (2015), comprising six putative populations of clonally propagated genotypes of different ecotypes (U: upland; L: lowland), ploidy levels (4X: tetraploid; 8X: octoploid) and geographical origins (S: South; W: West; N: North; E: East): U4X-N (135 plants), U8X-W (129 plants), U8X-E (97 plants), U8X-S (10 plants), L4X-NE (106 plants) and L4X-S (37 plants). These populations corresponded to 66 diverse accessions (Lu *et al.* 2013, Evans *et al.* 2015) with up to 10 individuals per accession.

In WS4U-C2, one individual was discarded so as to avoid assigning it to a population in AP, since it was too distantly related to the other individuals in BP (based on principal component analysis). In total, *n* = 760 individuals were considered in this analysis. The main goal of this study was to assess different methods for accommodating genetic heterogeneity when predicting phenotypic means in a given target population. Four targets were chosen, with a defined focus on tetraploid populations with at least 100 relatively homogeneous individuals: WS4U-C2 and Liberty-C2 (from BP), and U4X-N and L4X-NE (from AP).

### Marker data

Exome capture sequencing of individuals (parents in BP and clonally propagated plants in AP) was performed using the Roche-Nimblegen protocol for preparation of SeqCap EZ Developer libraries using the Roche-Nimblegen probeset ‘120911_Switchrass_GLBRC_R_EZ_HX1’ as described previously (Evans *et al.* 2014, 2015). Reads from sequencing were aligned to the hardmasked *P. virgatum* v1.1 reference genome (http://phytozome.jgi.doe.gov/pz/portal.html#!info?alias=Org_Pvirgatum). Counts of reads corresponding to alternate and reference alleles for each individual were then determined as described previously (for BP, Ramstein *et al.* 2016; for AP, Evans *et al.* 2014, 2015) at 2,179,164 single nucleotide polymorphism (SNP) loci, which were identified as polymorphic in two diversity panels: the Northern Switchgrass Panel, corresponding to AP (Evans *et al.* 2014, 2015), and a southern switchgrass panel (E.C. Brummer, unpublished). The numbers of alternate allele at the SNP loci were then called by using the expectation-maximization algorithm of Martin *et al.* (2010) fitted in each population (in BP) or accession (in AP) separately, under the assumption of disomic inheritance. Although this assumption is supported in switchgrass for tetraploid genotypes (Okada *et al*. 2010; Li *et al.* 2014), it does not hold for octoploid genotypes, which would presumably exhibit tetrasomic inheritance. However, we did not adapt the algorithm of Martin *et al.* (2010) to accommodate possible tetrasomic inheritance, as sequencing depth was deemed insufficient for calling intermediate heterozygotes (simplex and triplex) with high enough accuracy.

The resulting marker-data matrix consisted of expected allelic dosages (sums of alternate-allele counts weighted by their posterior probabilities, for every individual and SNP). The SNPs were then filtered based on the following criteria: (i) proportion of missing values strictly lower than 2%; (ii) minor allele frequency strictly greater than 1/2*n* and variance of expected allelic dosages strictly greater than 2(1/2*n*)(1 – 1/2*n*); (iii) *p*-value for Hardy-Weinberg equilibrium strictly greater than 10^−4^ in each BP population; (iv) availability of genomic-location information (as per v1.1 of the reference genome of *P. virgatum*). Missing values at SNPs were imputed by their mode in the whole sample. The resulting *n* ×; *m* filtered and imputed marker-data matrix **X** consisted of expected allelic dosages at *m* = 717,814 SNP markers.

### Phenotypic data

Populations in BP were assayed each year between 2012 and 2014, in Arlington, WI (USA), in a randomized complete block design, with four replicates for WS4U-C2 and three replicates for Liberty-C2. Populations in AP were assayed each year between 2009 and 2011 in Ithaca, NY (USA), in a sets-in-reps design, with two replicates per individual and 10 sets within each replicate, with each set comprising at most one individual from each of the 66 accessions in AP (Lu *et al.* 2013, Evans *et al.* 2015). In each panel, three phenotypic traits were considered: plant height, heading date and standability. Plant height (PH) was measured in centimeters as the height from the ground to the top of the tallest panicle. Heading date (HD) was measured in growing degrees days as the cumulated sum of daily average temperatures (in degrees Celsius; °C) above 10 °C, from January 1^s^^t^ to the day of heading, defined as the emergence of at least half of the panicles from the boot (Mitchell *et al.* 1997); daily average temperatures were estimated by the average of the minimum and maximum daily temperatures. Standability (St) was measured on a 0-10 scale to describe plants’ stature and stiffness, with 0 qualifying plants that are prostrate and 10 qualifying upright and rigid plants (Lipka *et al.* 2014).

Not all traits were measured every year in any given population: only HD was measured in all three years in AP populations and Liberty-C2. For all other cases, measurements were available for only a subset of years (Table 2).

**Table 2.**
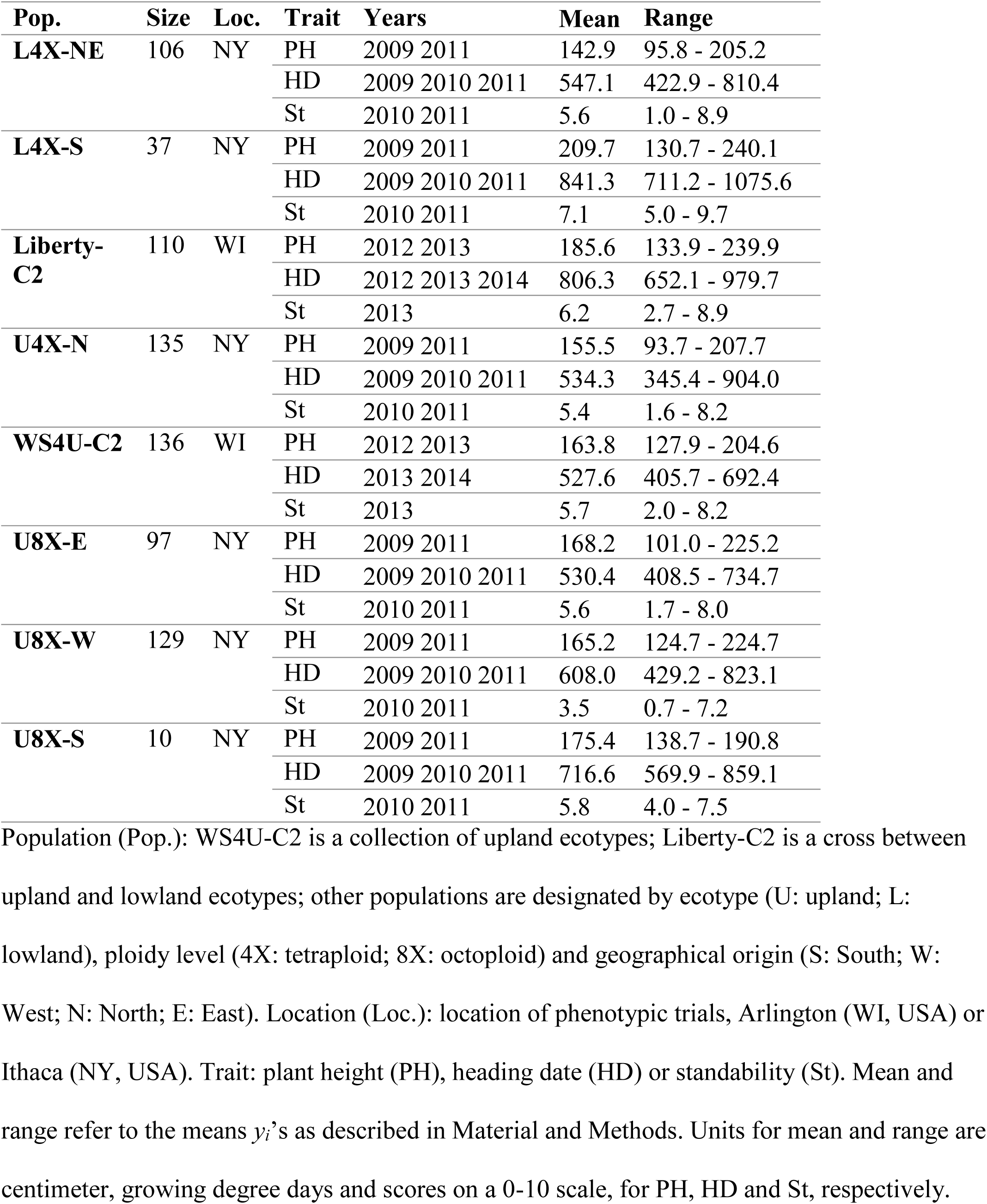
Description of populations and trait measurements

| **Pop.** | **Size** | **Loc.** | **Trait** | **Years** | **Mean** | **Range** |
| --- | --- | --- | --- | --- | --- | --- |
| **L4X-NE** | 106 | NY | PH | 2009 2011 | 142.9 | 95.8 - 205.2 |
|  |  |  | HD | 2009 2010 2011 | 547.1 | 422.9 - 810.4 |
|  |  |  | St | 2010 2011 | 5.6 | 1.0 - 8.9 |
| **L4X-S** | 37 | NY | PH | 2009 2011 | 209.7 | 130.7 - 240.1 |
|  |  |  | HD | 2009 2010 2011 | 841.3 | 711.2 - 1075.6 |
|  |  |  | St | 2010 2011 | 7.1 | 5.0 - 9.7 |
| **Liberty-C2** | 110 | WI | PH | 2012 2013 | 185.6 | 133.9 - 239.9 |
|  |  |  | HD | 2012 2013 2014 | 806.3 | 652.1 - 979.7 |
|  |  |  | St | 2013 | 6.2 | 2.7 - 8.9 |
| **U4X-N** | 135 | NY | PH | 2009 2011 | 155.5 | 93.7 - 207.7 |
|  |  |  | HD | 2009 2010 2011 | 534.3 | 345.4 - 904.0 |
|  |  |  | St | 2010 2011 | 5.4 | 1.6 - 8.2 |
| **WS4U-C2** | 136 | WI | PH | 2012 2013 | 163.8 | 127.9 - 204.6 |
|  |  |  | HD | 2013 2014 | 527.6 | 405.7 - 692.4 |
|  |  |  | St | 2013 | 5.7 | 2.0 - 8.2 |
| **U8X-E** | 97 | NY | PH | 2009 2011 | 168.2 | 101.0 - 225.2 |
|  |  |  | HD | 2009 2010 2011 | 530.4 | 408.5 - 734.7 |
|  |  |  | St | 2010 2011 | 5.6 | 1.7 - 8.0 |
| **U8X-W** | 129 | NY | PH | 2009 2011 | 165.2 | 124.7 - 224.7 |
|  |  |  | HD | 2009 2010 2011 | 608.0 | 429.2 - 823.1 |
|  |  |  | St | 2010 2011 | 3.5 | 0.7 - 7.2 |
| **U8X-S** | 10 | NY | PH | 2009 2011 | 175.4 | 138.7 - 190.8 |
|  |  |  | HD | 2009 2010 2011 | 716.6 | 569.9 - 859.1 |
|  |  |  | St | 2010 2011 | 5.8 | 4.0 - 7.5 |

Population (Pop.): WS4U-C2 is a collection of upland ecotypes; Liberty-C2 is a cross between
upland and lowland ecotypes; other populations are designated by ecotype (U: upland; L:
lowland), ploidy level (4X: tetraploid; 8X: octoploid) and geographical origin (S: South; W:
West; N: North; E: East). Location (Loc.): location of phenotypic trials, Arlington (WI, USA) or
Ithaca (NY, USA). Trait: plant height (PH), heading date (HD) or standability (St). Mean and
range refer to the means $y_i$ 's as described in Material and Methods. Units for mean and range are
centimeter, growing degree days and scores on a 0-10 scale, for PH, HD and St, respectively.

In BP, observational units were plants within half-sib families from a given genotype (maternal parent) *i*. Half-sib families were arranged in a randomized complete block design and assayed in multiple years; so the following model was fitted to phenotypic measurements *P*_*ijkl*_, to estimate half-sib family effects *f*_*i*_’s:

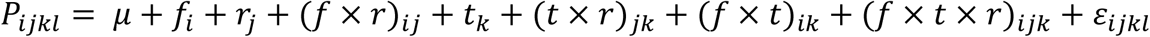

where *μ* is the population mean; *f*_*i*_, *r*_*j*_ and *t*_*k*_ are the effects of half-sib family *i* (fixed), block *j* (random) and year *k* (random) respectively; ×; indicates interactions (random); *ε*_*ijkl*_ are residuals for plant *l* within plot *ij* in year *k*.

In AP, observational units were clones of a given genotype *i*. Genotypes were arranged in a sets-in-reps design and assayed in multiple years; so the following model was fitted to measurements *P*_*ijkl*_ to estimate genotype effects *g*_*i*_’s:

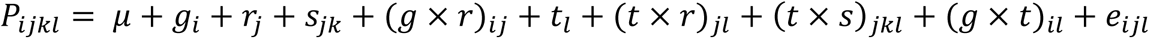

where *μ* is the panel mean; *g*_*i*_, *r*_*j*_, *s*_*jk*_ and *t*_*l*_ are the effects of genotype *i* (fixed), replicate *j* (random), set *k* within replicate *j* (random) and year *l* (random) respectively; ×; indicates interactions (random); *e*_*ijl*_ are residuals for clone *ij* in year *l*.

The models described above conform to analyses of strip-plot (split-block) designs (Steel *et al.* 1996), in which years and genotype classes (half-sib families in BP, individual genotypes in AP) are whole-plot factors in cross-classification, in a given replicate; sub-plot factors are combinations of years and genotype classes. For each random term, the corresponding effects were modelled as independent and identically normally distributed. The linear mixed models described above were fitted using ASREML-R (Butler *et al.* 2009).

Effects *f*_*i*_’s are transmitting abilities of genotypes, so that 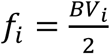, where *BV*_*i*_ is the breeding value of genotype *i*. In comparison, effects *g*_*i*_’s are genotypic values, such that *g*_*i*_ = *BV*_*i*_ + *Δ*_*i*_, where *Δ*_*i*_ is the deviation from additivity due to dominance and/or epistasis. Outcomes of interest for genomic prediction were set to be genotype means *y*_*i*_’s such that 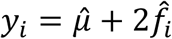 in BP and 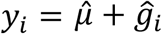 in AP.

### Population structure data

#### Admixture analysis

The soft clustering model from the ADMIXTURE software was fitted on the whole sample and the whole set of SNPs, i.e., without selection on individuals or markers (Alexander *et al.* 2009). Based on the 10-fold cross-validation implemented in ADMIXTURE (Alexander *et al.* 2011), the number of population clusters in the admixture model was set to *K* = 7, as cross-validation error reached a plateau at that value (Figure S1). The resulting *n* ×; *K* matrix **A** of admixture coefficients comprised inferred membership probabilities at each cluster (Figure 1a). For convenience (in prediction models), minimum values in **A** (10^−5^) were set to zero while ensuring that each row still summed to one.

**Figure 1.**
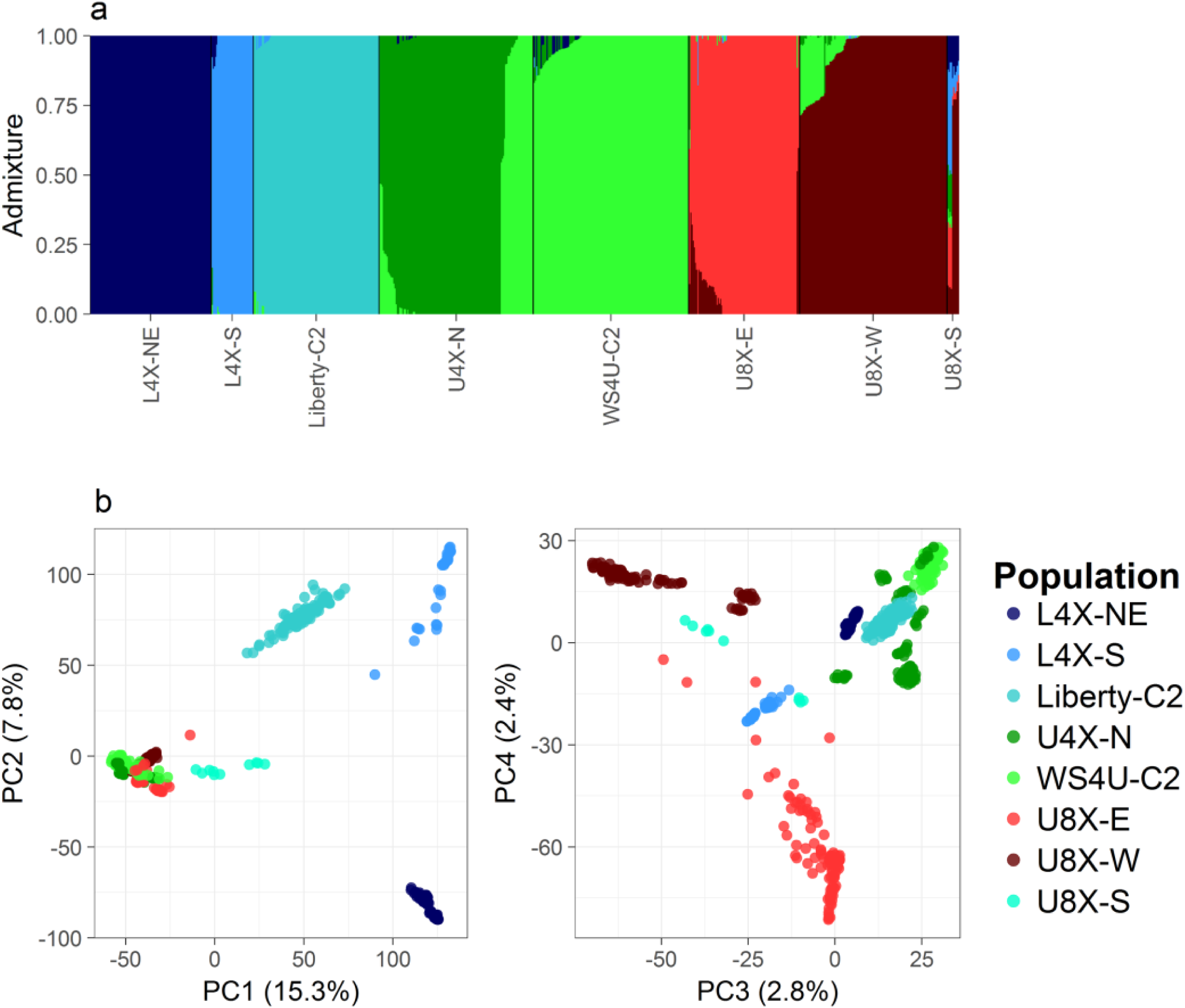
Population structure in the sample (a) Admixture plot of the whole sample, with colors designating the seven inferred population clusters, which roughly matched populations, with the exception of U8X-S which displayed strong admixture; (b) Principal component analysis (PCA) plot of the whole sample of 760 individuals, with colors designating the eight populations.

#### Principal component analysis

Principal component analysis (PCA) was performed on the whole sample and the whole set of SNPs. The number of principal components (PCs) to choose for depicting population structure was chosen based on the proportion of variance explained and the grouping patterns captured by PCs (Figure 1b). The resulting *n* ×; *d* PC matrix **P** consisted of coordinates for each individual at the first *d* = 4 PCs.

### Genomic prediction models

All linear mixed models described below were fitted using the R package rrBLUP (Endelman 2011).

For a given marker-data matrix **X** and vector **y** of outcomes, the standard ridge regression BLUP model (RR-BLUP; BLUP: best linear unbiased predictor) is described as follows:

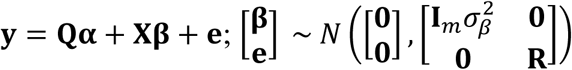

where **y** is the *n*-vector of outcomes (*y*_*i*_’s, as described above); **X** is the *n* ×; *m* marker-data matrix (consisted of the expected allelic dosages, here with no transformation or scaling), and **β** is the *m*-vector of marker effects, assumed independent with variance 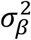 (***I***_*m*_ is the *m* ×; *m* identity matrix); **Q** is the *n* ×; *p* matrix depicting the population mean structure in the sample, and **α** is the *p*-vector of associated effects; **R** is the covariance matrix of errors, possibly accommodating correlations and differences in variance (heteroscedasticity) among errors. Often, errors are considered to be independent and identically distributed, such that 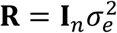, with *I*_*n*_ the *n* ×; *n* identity matrix and 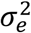 the error variance.

Let **u** = **Xβ**, so that Var 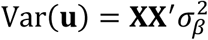, by identical mean structure **Qα** and variance structure Var 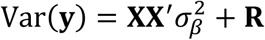 (Henderson, 1984), the RR-BLUP model is equivalent to the following genomic BLUP (GBLUP) model:

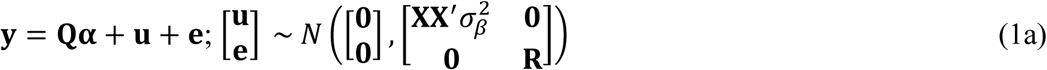

In the RR-BLUP model, regressing out the mean-structure matrix **Q** from **X** yields the following equivalent model, where mean- and variance-structure matrices are orthogonal, i.e. columns from one matrix to another are now uncorrelated (see Appendix A1 for a general proof):

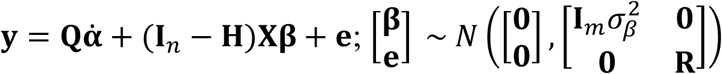

where **H = Q(Q^′^*R*^−1^Q)^−1^Q^′^*R*^−1^** is the matrix of projection onto the column space of **Q**;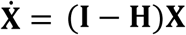 is the adjusted matrix of (residual) marker variables, made orthogonal to 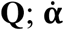 is the new vector of fixed effects. With **Q** = **1**_***n***_ (**1**_***n***_ is a *n*-vector of ones) and ***R*** = ***I***_*n*_, the mean structure 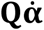 is simply an intercept and 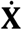 is the matrix of marker variables centered around their respective mean, as often used in genomic prediction studies (Hayes *et al.* 2009b, de los Campos *et al.* 2013). With general **Q**, the mean structure 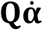 is an individual-specific mean for **y** with respect to the specific population membership of each individual and 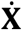 is the matrix of marker variables centered around their respective individual-specific means.

By identical mean and variance structures, the previous model is equivalent to the following alternate GBLUP model:

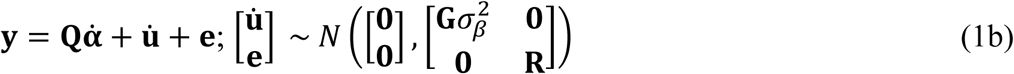

where 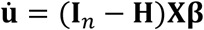 is the *n*-vector of genomic breeding values centered around population means 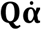, and 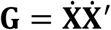 is the genomic relationship matrix for 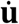 (here unscaled).

Following the recommendations of Phocas and Laloë (2004), we chose to simply use **Q** = **1**_***n***_ to define the mean structure in all fitted models. Also, we chose not to model heteroscedasticity of errors and used 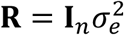. Therefore, the covariance matrix 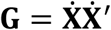 was simply proportional to the standard genomic relationship matrix 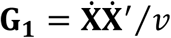 of VanRaden (2008), where 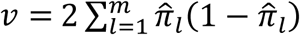 is a scaling factor depending on estimated allele frequencies 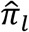’*s*. Notably, the matrix **G** derived here will account for correlations and heteroscedasticity of errors, whenever **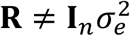** (the projector **H** is a function of **R**^−1^). To our knowledge, the matrix **G1** has typically been used in GBLUP models, even when 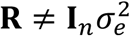 as in weighted GBLUP models on deregressed proofs.

### Optimization methods

Hereafter, the testing set TS is defined as the set of individuals left out for model validation. The calibration set CS is the set of individuals used to fit the prediction models, which excludes TS but does not necessarily consists of all remaining (available) individuals.

In this study, we adapted model (1a) or (1b) to four different approaches: (i) the control procedure, consisting in including in the CS only individuals from the same target group as the TS; (ii) single-population models, consisting in fitting a GBLUP model to all available individuals, with possibly some regularization on genomic relationships; (iii) instance selection, consisting in including a subset of available individuals in the CS, so as to optimize some selection criterion; and (iv) multi-population models, consisting in modelling population heterogeneity for the outcome on all available individuals, based on population structure data (PCs in **P** or admixture coefficients in **A**).

#### Control procedure (Target)

In the control procedure (*Target*), we fitted model (1a), with the CS restricted to individuals belonging to the same population as the TS. This method corresponds to a typical choice of relying only on individuals that have genetic architectures that are *a priori* similar to those in the TS.

#### Single population models (SPM-GRM, SPM-GLASSO)

Here, single-population models (*SPM*) are defined as the basic models incorporating information from all available individuals, with no modelling of population heterogeneity. The following general model was fitted:

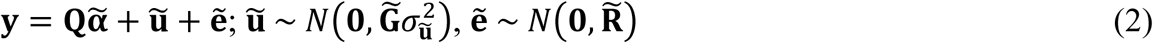

where 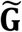 is some matrix depicting relationship among breeding values 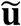.

We considered two types of matrices for 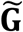: the original unscaled genomic relationship matrix 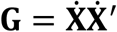 (*SPM-GRM*) and a regularized form of **G**, where relationships are shrunk for potentially higher estimation accuracy of relationships (*SPM-GLASSO*).

In *SPM-GRM* (the reference *SPM* procedure), since 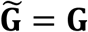, model (2) was equivalent to model (1b), so this approach simply corresponded to fitting a GBLUP model to all available individuals.

In *SPM-GLASSO* (the proposed *SPM* procedure), following Fan *et al.* (2013), 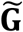 was decomposed as 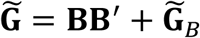, where **B** consisted of the first *t* PCs of **G** and 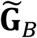*B* was a regularized form of **G**_*B*_ = **G** – **BB^′^**. Matrix **BB^′^**is the dense part of the relationship matrix **G**, representing resemblance among individuals through common structural factors. Here this matrix depicted relationships at the population level, through the *t* leading PCs of **G**. In contrast, matrix **G**_*B*_ represented (recent) relationships conditional on population structure, similarly to the adjusted relationships introduced by Thornton *et al.* (2012) and Conomos *et al.* (2016), with the difference that here coefficients in **G**_*B*_ are not scaled for direct estimation of recent-kinship coefficients. In principle, most coefficients in **G**_*B*_ should be close to zero, as there should exist only few familial relationships within the sample. So **G**_*B*_ may be assumed to be sparse. Fan et *al.* (2013) suggested that matrix **G**_*B*_ be regularized by adaptive thresholding (Cai and Liu 2011). However, we chose to perform regularization by the graphical LASSO (Friedman *et al.* 2008) so as to shrink coefficients in **G**_*B*_ while inferring a sparse precision (inverse covariance) matrix 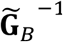, which yielded a sparse graph of relationships among individuals (a zero *ij*-element in 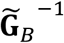 indicates that individuals *i* and *j* are unrelated conditionally on all other individuals, which corresponds to no edge between nodes *i* and *j* in the underlying graph of recent relationships).

The graphical LASSO infers a sparse precision matrix **Σ**^−1^ by maximizing the Gaussian likelihood of the data, penalized by a *L*_*1*_-norm penalty *λ*‖**Σ**^−1^‖1, where *λ* is the regularization parameter and ‖**Σ**^−1^‖_1_ is the sum of absolute values in **Σ**^−1^ (Friedman *et al.* 2008).

Regularization of **G** was performed as follows:

1. Performing eigenvalue decomposition on **G** to obtain **B** and decompose **G** into **BB**^′^+ **G**_*B*_
2. Standardizing **G**_*B*_ to obtain the corresponding correlation matrix **Γ**_*B*_: **Γ**_*B*_ = diag(**G**_*B*_)^−1/2^**G**_*B*_diag(**G**_*B*_)^−1/2^
3. Applying the graphical LASSO algorithm to **Γ**_*B*_, to obtain the regularized correlation matrix 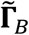
4. Rescaling **Γ**_*B*_ to obtain 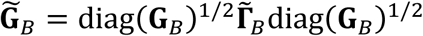 and then 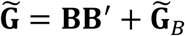

The graphical LASSO algorithm was run using the R package huge (Zhao *et al.* 2012). The regularization parameter *λ* was determined so as to maximize the restricted maximum likelihood (REML) of model (2) in a grid search, with *λ* being the *q*-quantile of absolute values in **Γ**_*B*_ and *q* varying from 0.05 to 1 by step of 0.05.

With **Q** = **1**_***n***_, the number *t* of PCs in **B** was set to *d* = 4 (*d* is the number of PCs chosen to reflect population structure in **P**). However, when **Q** actually depicts population structure, e.g. when **Q** = [**1**_***n***_ **P**] or **Q** = **A**, the matrix **G** already reflects relationships among individuals conditionally on population structure (so **G** should already be sparse), and *t* may simply be set to zero (i.e., regularization may be performed on **G** directly). Notably, when 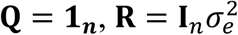 and the CS consists of the whole sample, **B** simply equals **P**.

#### Instance selection (IS-Rel, IS-QP)

In instance selection (*IS*), we fitted model (1a) on a subset of all available individuals. We selected individuals deterministically (i.e., without using random searches through possible calibration sets) by first including individuals with highest scores (as defined below), so as to optimize a selection criterion. We chose to maximize the mean coefficient of determination CD_mean_ (Laloë 1993) for the TS, with no contrast so that this selection criterion simply corresponded to the model-based estimate of the mean squared prediction accuracy (reliability) with respect to **u** in the TS, i.e. 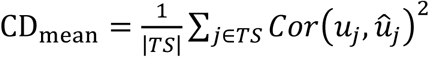, with |*TS*| the size of the TS, 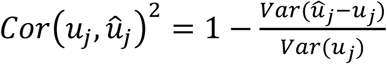, where 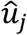 is the BLUP of *u* j, *Var*(*ûj* – *u*j) and *Var*(*u*j) are inferred from the model fit (Searle *et al.* 2006).

We considered two types of scores (*w*_*i*_’s), for two different procedures: *IS-Rel* and *IS-QP*.

In *IS-Rel* (the reference *IS* procedure), 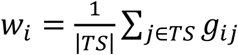, with *g*_*ij*_ being the *ij*-element of 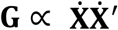. So *w*_*i*_ simply reflected the average relationship between individual *i* and the TS.

In *IS-QP* (the proposed *IS* procedure), we inferred the scores *w*_*i*_’s on all available individuals, so as to minimize the difference between the average genotype in the TS and the weighted average of genotypes in remaining individuals (TS^C^), with weights *w*_*i*_’s. Formally, let 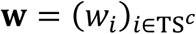 such that *w*_*i*_ ≥ 0 for all *i* ∈ TS^*c*^ and **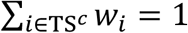**, we minimized 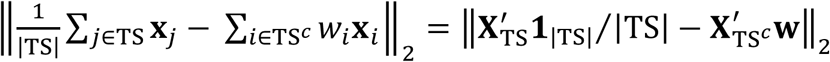, with ‖. ‖_2_ being the Euclidean norm, and subscripts referring to subsets on rows in vectors or matrices (**x**_*i*_ refers to the *m*-vector of marker variables for individual *i*). Equivalently, we minimized 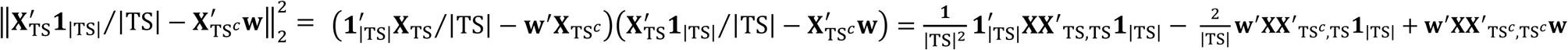. Since the first term in the last sum is constant with respect to *w*_*i*_’s, the optimal **w** solved the following quadratic programming (QP) problem: minimizing 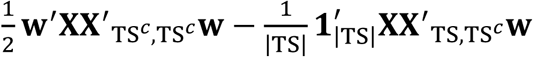 subject to *w*_*i*_ ≥ 0 for all *i* ∈ TS^*c*^ and 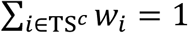. This problem is similar to the general QP problem for feature selection introduced by Rodriguez-Lujan *et al.* (2010), i.e., minimizing 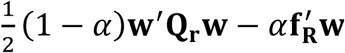, subject to *w*_*i*_ ≥ 0 for all *i* and **Σ**_*i*_ *w*_*i*_ = 1, where vector **f**_**R**_ measures relevance of features with respect to a given outcome; matrix **Q**_**r**_ measures the redundancy among features; and *α* ∈ [0,1] sets the relative importance of each term in the sum. The QP problem could have been defined freely, but our initial motivation allowed us to naturally set 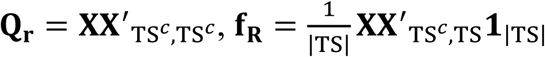 and 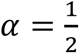. Compared to *IS-Rel, IS-QP* has two advantages: a solution **w** will represent a compromise between relevance (average of relationships 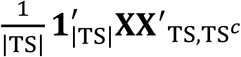 from TS^*c*^ to TS) and redundancy (relationships 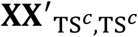 within TS^*c*^), so scores from *IS-QP* should favor more diverse sets of individuals compared to *IS-Rel*; also, relationships used are not adjusted by allele frequencies (equivalently, they do not depend on any projector **H**), so they could be less prone to misrepresentations of relationships, through inappropriate centering of marker variables.

The QP problem in *IS-QP* was solved using the R package quadprog (https://CRAN.R-project.org/package=quadprog). In optimization, we considered selecting 5% to 100% of individuals in CS, by step of 5%, selecting the subset of individuals which maximized CD_mean_.

The prediction accuracy of *IS* methods was assessed with free CS sizes (i.e., as optimization procedures), but was also assessed with fixed CS sizes, with no selection of subset based on CD_mean_: for a given CS size |CS|, only the first |CS| individuals with the highest scores *w*_*i*_’s were included in the CS. Such assessments were intended to reflect the usefulness of *IS* procedures in conditions of limited resources for phenotyping (and calibration of prediction models). In this context, *IS-Rel* and *IS-QP* were compared to random selection, which simply consisted in randomly selecting |CS| individuals for model fitting. For a given combination of TS, |CS| and outcome, the accuracy from random selection was the average of accuracies over 200 random draws. Each draw corresponded to one random attribution of scores (*w*_*i*_’s) to individuals.

#### Mixed population models (MPM-Mixture, MPM-Matérn)

Mixed-population models (*MPM*) were extensions of model (1a) intended to accommodate population heterogeneity. The following general model was fitted:

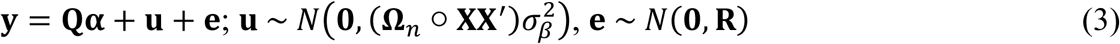

where ○ is the element-wise (Hadamard) product, and **Ω**_*n*_ is a *n* x *n* covariance matrix depicting population differentiation among individuals (see Appendix A2 for derivations and technical details). To parsimoniously estimate **Ω**_*n*_, we used two different procedures: *MPM-Mixture* (based on **A**) and *MPM-Matérn* (based on **P**). In both procedures, we did not model any heteroscedasticity for additive genetic effects **u**.

In *MPM-Mixture* (the reference *MPM* procedure), **Ω**_*n*_ = ρ**AΘ**_*K*_**A**^′^+ (1 – ρ)**J**_*n*_, where **J**_*n*_ is the *n* x *n* matrix of ones and **Θ**_*K*_ is a *K* x *K* matrix depicting relationships among population clusters as inferred in **A**. Here, we simply set **Θ**_*K*_ = **I**_*K*_ (**I**_*K*_ is the *K* x *K* identity matrix), so **Ω**_*n*_ = ρ**AA**^′^+ (1 – ρ)**J**_*n*_. Therefore in this procedure, ρ ∈ [0,1] set a trade-off between the case where relationships were cluster-specific (ρ = 1) and the case where relationships assumed one single homogeneous population for all individuals (ρ = 0). This approach is similar (but not exactly equivalent) to the *K-*kernel method of Heslot and Jannink (2015), which considered a similar balance between cluster-specific and overall relationships, but using **G**_1_ for relationships (VanRaden 2008), instead of **XX**^′^, and considering only discrete population clusters (in which case values in **A** would then be only 0 or 1). Alternatively, *MPM-Mixture* may be viewed as a multi-kernel model where 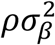 and 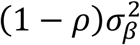 are the variances components respectively associated to cluster-specific and main marker effects.

In *MPM-Matérn* (the proposed *MPM* procedure), **Ω**_*n*_ = (*k*_*v,h*_(***p***_*i*_, ***p***_*j*_))_*n*×;*n’*_, where *k*_*v,h*_ is a Matérn kernel function of **p**_*i*_ and 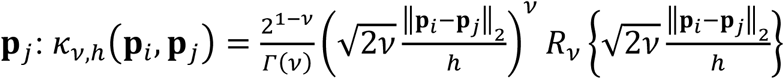, ‖**p**_*i*_ – **p**_*j*_‖_2_ is the Euclidean distance between the *d*-vectors of PC coordinates for any pair (*i, j*) of individuals, *v* > 0 is a shape parameter, *h* > 0 is a scale parameter, and *R*_*v*_{.} is the modified Bessel function of the second kind, of order *ν* (Abramowitz and Stegun 1984, Ober *et al.* 2011). Matérn functions have been used in various contexts, including in genomic prediction for depicting relationships among individuals (Ober *et al.* 2011). Here, we used Matérn functions to depict relationships among populations, with the input ‖**p**_*i*_ – **p**_*j*_‖_2_ representing differentiation with respect to population structure in *d* = 4 orthogonal directions. We used Matérn functions instead of more typical kernel functions (e.g., an exponential or Gaussian kernel function) to allow for some flexibility in the shape of the correlation in **Ω**_*n*_: *v* = 0.5 and *v* = ∞ correspond respectively to the exponential and Gaussian kernels as special cases, while different shapes can also be fitted (Ober *et al.* 2011).

The parameter *ρ* in *MPM-Mixture* was estimated by maximizing the restricted likelihood of model (3) using the optimization algorithm implemented in the R function *optimize*. The parameters *ν* and *h* in *MPM-Matérn* were estimated by maximizing the restricted likelihood of model (3) using the Nelder-Mead algorithm implemented in the R function *constrOptim*, with constraints for positivity. In order to control (to some extent) for the possible presence of local maxima in the restricted likelihood surface in *MPM-Matérn*, we used four different starting points (*v*_0_, *h*_0_): (0.5, *D*_*max*_/2), (0.5, *D*_*max*_), (10, *D*_*max*_/2) and (10, *D*_*max*_), with *D*_*max*_the maximum distance ‖**p**_*i*_ – **p**_*j*_‖_2_ observed over pairs of individuals (*i, j*).

### Validations

We assessed the accuracy of our prediction procedures by cross-validation (CV): for each target (L4X-NE, U4X-N, Liberty-C2 or WS4U-C2), we used as the TS a random subset of the target sample. The size of the TS was one fifth of the target sample size. All remaining individuals were used as input to the prediction procedures (*Target, IS, SPM* or *MPM*), with some CS selection in *Target* and *IS*. Such validations were replicated *n*_*rep*_ = 20 times for each target.

Prediction procedures were evaluated for accuracy by 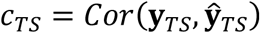, i.e., the correlation between “observed” and predicted outcomes in a given TS. To assess the significance of differences in prediction accuracy between two procedures, we performed a t-test on 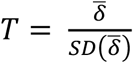 where 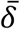 is the average of **δ** = *Z*(**c**_*t*_) – *Z*(**c**_0_); **c**_*t*_ (**c**_0_) is the vector of prediction accuracies over testing sets for the tested procedure (baseline procedure); and *Z* is the Fisher transformation. The standard error of the mean difference in prediction accuracy, 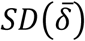, was estimated in two different ways: (liberal t-test) 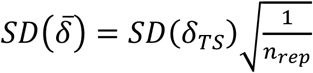 where *SD*(δ_*TS*_) is the standard deviation of **δ**, with all testing sets assumed to be independent datasets; (conservative t-test) based on the first method of Nadeau and Bengio (2003), 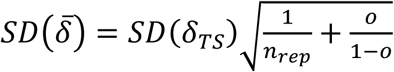, where redundancy over testing sets is accounted for by the additional term 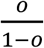, with *o* being the expected fraction of overlap among testing sets; here *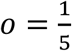* and *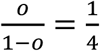* because testing sets were random subsets consisting of a fifth of any given target sample. We considered that this approach for estimating (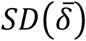) was conservative because Nadeau and Bengio (2003) derived it by assuming that the CV criterion (the “loss function”, analog here to *Z*(*c*_*TS,t*_) – *Z*(*c*_*TS*,0_), for a given TS) did not depend on the CS instances, given a particular CS size. Therefore the adjustment from Nadeau and Bengio (2003) may have overestimated the correlation among values of the CV criterion across replicates, since prediction procedures are probably quite sensitive to differences in the composition of the CS. Furthermore, some procedures actually differed in CS size (*Target, IS*, and *SPM*/*MPM*). In all comparisons between procedures, we reported the results from both tests in order to characterize the significance of differences in prediction accuracy.

## Data availability

Population information (population assignment and geographical origin of genotypes, when available), raw phenotypic data (trait measurements at individual plants) and estimated genotype means (for maternal parents in BP and individuals in AP) are available in Files S1, S2 and S3, respectively. These supplementary files as well as the marker data (expected allelic dosages at the 717,814 selected SNP markers; in .rds format readable in R) can be downloaded from dfrc.wisc.edu/sniper.

## RESULTS

### Population structure in the sample

Seven population clusters were inferred from the ADMIXTURE software (Figure S1; Alexander *et al.* 2009). These clusters corresponded roughly to populations L4X-NE, L4X-S, Liberty-C2 and U4X-N, WS4U-C2, U8X-E, U8X-W. One population with little representation in our sample, U8X-S, appeared to be of mixed origin (Figure 1a). The other populations generally displayed a low level of admixture, with relatively few individuals having intermediate admixture coefficients. There seemed to be some admixture involving upland populations (WS4U-C2 and U4X-N, WS4U-C2 and U8X-W, U8X-E and U8X-W), with even some shared ancestry between WS4U-C2 and U4X-N. The PCA confirmed that population structure was relatively discrete (Figure 1b). Unsurprisingly, the first PCs separated genotypes by ecotype while the second PC reflected geographical origin within the lowland ecotype (Lu *et al.* 2013, Evans *et al.* 2015). The third and four PCs discriminated upland genotypes by geographical origin and ploidy level, and distinguished L4X-S from the two other lowland populations (L4X-NE and Liberty-C2).

Differences in mean and range among populations were quite typical of previously reported differences between ecotypes (Table 2; Casler 2012). Indeed, L4X-S and Liberty-C2 (populations of lowland origin) had high mean values and range values for PH, HD and St, compared to upland populations (excluding U8X-S). However, L4X-NE stood out as a lowland population for being relatively short, early-flowering, and prone to lodging, with corresponding values for PH, HD, and St more similar to those of the upland populations.

### Single-population models and relationships in the sample

Here, marginal genomic relationships were defined as the elements of 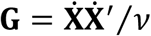, with 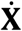 consisting of centered marker variables, and *?* being some scaling factor. The strong and quite discrete population structure in the sample translated into multimodal marginal genomic relationship coefficients, with the multiple peaks in off-diagonal elements of **G** reflecting differentiation of population with respect to allele frequencies (Figure 2a). Conditioning relationships on population structure (as depicted by the first four PCs of 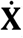) yielded the matrix **G**_*B*_, with **G**_*B*_ = **G** – **BB**^′^and **B** consisting of the PCs chosen to reflect structure in **G** (Fan *et al.* 2013). The conditional genomic relationships seemed sparser, in the sense that they appeared to cluster around zero, so most individuals could be assumed to be unrelated after accounting for population structure in the sample. In our particular study, conditional relationships in **G**_*B*_ were all the more relevant that among-population variation, captured by **BB**^′^, contributed little to variation within any given TS, because a TS generally consisted of selection candidates from a relatively homogeneous target sample (made of individuals from WS4U-C2, Liberty-C2, U4X-N or L4X-NE).

**Figure 2.**
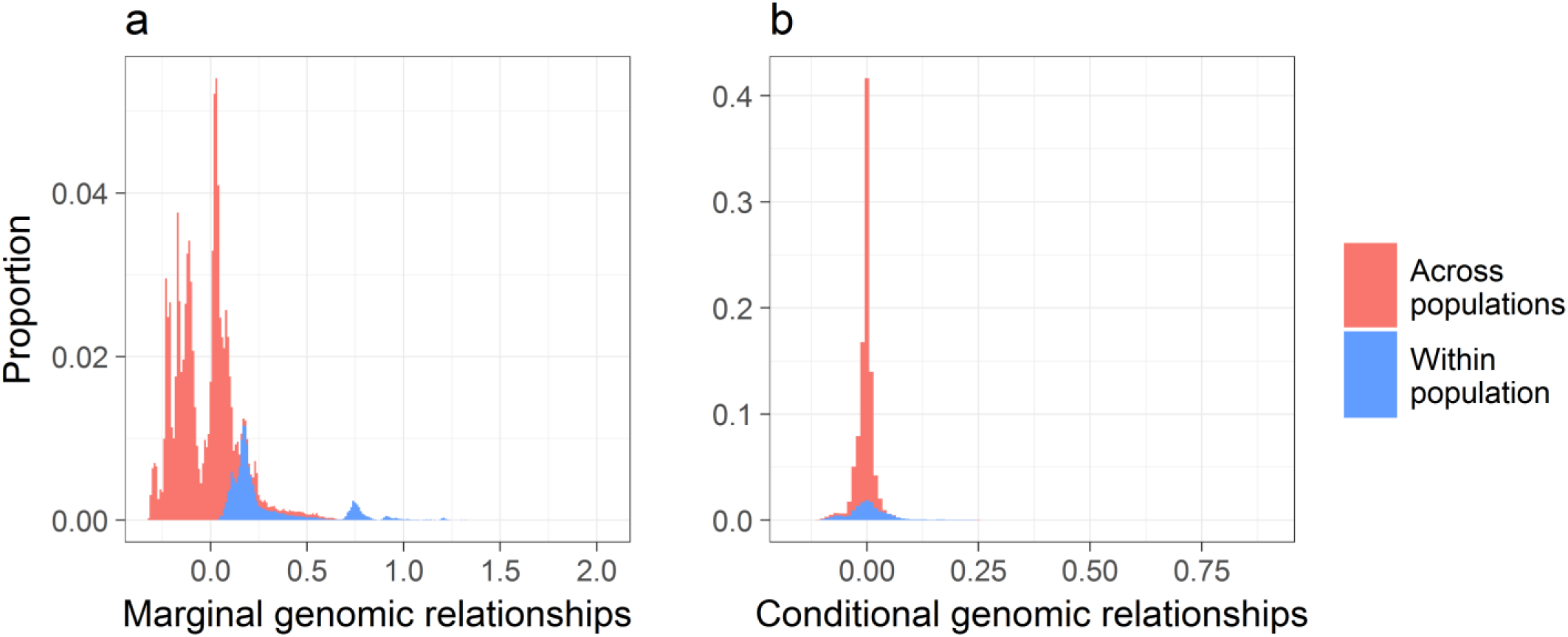
Histograms of genomic relationships in the sample (a) Marginal genomic relationships, scaled as in VanRaden (2008): off-diagonal elements of **G**/*v*, with 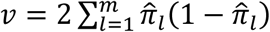 (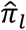: estimated allele frequency at marker *l*). (b) Genomic relationships conditional on population structure, as captured by PCs, also scaled as in VanRaden (2008): off-diagonal elements of **G**_B_/*v*, with **G**_B_ = **G** – **BB**^′^(**B** consists of the first d = 4 PCs of **G**).

The *SPM-GLASSO* model did not yield substantial increases in quality of fit, compared to *SPM-GRM*, when fitted either to the whole sample or to calibration sets in cross-validation (CV) (Table 3). However, it is unclear to what extent a substantial improvement in fit should be expected from *SPM-GLASSO*: on the one hand, *SPM-GLASSO* relies on one additional parameter *λ* compared to *SPM-GRM*, but on the other hand this parameter results in less complex relationships within 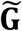 compared to **G** (Foygel and Drton 2010). In general, quite small regularization parameters (*λ*) were selected based on REML: when fitted to the whole sample, *SPM-GLASSO* selected values of *λ* between 0.002 (for PH) and 0.007 (for St), corresponding respectively to 0.20- and 0.45-quantiles of absolute correlations from **G**_*B*_. As a result, the inferred graphs were rather dense, with average degrees (number of neighbors by node/individual in the graph) ranging from 217 to 458 (Figure 3). However, even at such low regularization levels, some noticeable features of populations emerged from the inferred graphs (Figure 3): WS4U-C2, U4X-N and U8X-E appeared quite connected to one another; U8X-W also showed some connection with other upland populations but seemed more distinct, as reflected by a relatively lower average degree (Figure S2); Liberty-C2 and L4X-S were somewhat connected to both upland and lowland populations, which certainly explains why their individual degrees were generally high (Figure S2); most notably, L4X-NE displayed an outstandingly low level of connection with the other populations, which translated in a clear separation of this population in the graph, after placing the nodes based on a force-directed algorithm (Fruchterman and Reingold 1991). These features exemplify the usefulness of conditional relationships and their associated graphs for describing relationships among individuals and, potentially, serving as input to other types of procedures, e.g., instance selection.

**Table 3.** Parameter estimates, likelihood-ratio test statistic and *p*-value, by trait and procedure

| Trait | Procedure | Parameter estimate | LRT statistic | LRT $p$-value |
| --- | --- | --- | --- | --- |
| **PH** | *SPM-GLASSO* | $\lambda$: 0.002 (0.001-0.002) | 0.76 (0.37-1.70) | 0.38 (0.19-0.55) |
| | *MPM-Mixture* | $\rho$: 0.000 (0.000-0.009) | 0.00 (0.00-0.05) | 1.0 (0.82-1.0) |
| | *MPM-Matérn* | $h^*$: 0.525 (0.221-0.580)<br>$\nu$: 0.625 (0.550-0.884) | 7.39 (4.54-9.94) | 0.025 (0.0069-0.10) |
| **HD** | *SPM-GLASSO* | $\lambda$: 0.003 (0.002-0.003) | 2.29 (1.09-3.76) | 0.13 (0.052-0.30) |
| | *MPM-Mixture* | $\rho$: 0.434 (0.354-0.569) | 15.89 (13.85-18.82) | $6.7 \times 10^{-5}$ ($1.4 \times 10^{-5}$-$2.0 \times 10^{-4}$) |
| | *MPM-Matérn* | $h^*$: 0.325 (0.275-0.500)<br>$\nu$: 0.619 (0.537-0.761) | 42.76 (32.68-46.88) | $5.2 \times 10^{-10}$ ($6.6 \times 10^{-11}$-$8.0 \times 10^{-8}$) |
| **St** | *SPM-GLASSO* | $\lambda$: 0.007 (0.000-0.019) | 0.56 (0.00-2.46) | 0.45 (0.12-1.0) |
| | *MPM-Mixture* | $\rho$: 0.138 (0.115-0.159) | 5.72 (4.42-7.07) | 0.017 (0.0078-0.036) |
| | *MPM-Matérn* | $h^*$: 0.134 (0.125-1.150)<br>$\nu$: 9.049 (0.650-10.014) | 7.59 (5.55-8.54) | 0.022 (0.014-0.062) |

In parentheses: range over cross-validation replicates and target populations. Trait: plant height (PH), heading date (HD) or standability (St). *SPM-GLASSO*: Single-population model based on regularized relationships ( $\lambda$ : regularization parameter); *MPM*: Multi-population model with among-population correlations based on admixture coefficients (*MPM-Mixture*; $\rho$ : Mixture parameter) or PC distances (*MPM-Matérn*; $\nu$ : shape parameter; $h^* = h/D_{max}$ , with $h$ the scale parameter and $D_{max}$ , the maximum distance $\|\mathbf{p}_i - \mathbf{p}_j\|_2$ observed over pairs of individuals). LRT (likelihood-ratio test) statistic: $-2\log(L_0/L_1)$ where $L_0$ is the REML of *SPM-GRM* (GBLUP) and $L_1$ is the REML of one of the procedure described here; $p$ -values were obtained from a $\chi^2$ -distribution with one (*SPM-GLASSO*, *MPM-Mixture*) or two (*MPM-Matérn*) degrees of freedom.

**Figure 3.**
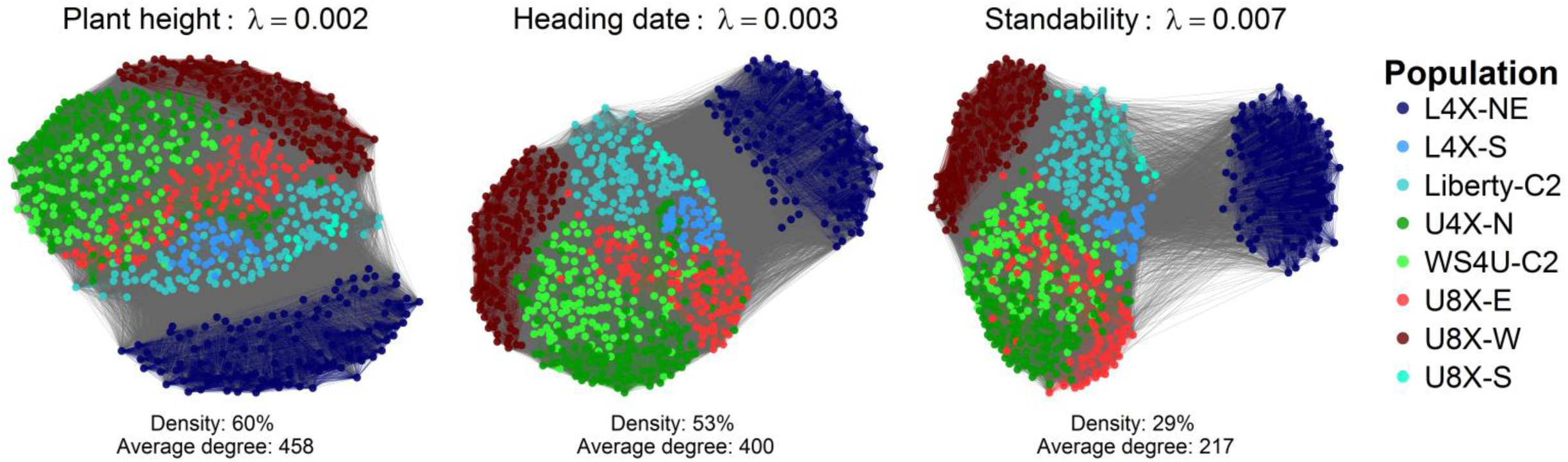
Inferred graphs of relationships, conditional on population structure Each graph represents the relationships as depicted from 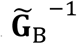 for a given trait, where 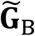 is the regularized matrix of conditional relationships obtained by the graphical LASSO applied to the whole sample of individuals (the absence of edge between two individuals is indicated by a zero *ij*-element in 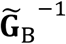). Nodes (individuals) were positioned using the force-directed placement algorithm of Fruchterman and Reingold (1991), as implemented in function ggnet (R package GGally), so aggregation of nodes reflects connectedness.

For prediction in a given TS, the control procedure (*Target*) consisted in restricting the CS to the subset of the sample belonging to the same population as the TS. Compared to *Target, SPM-GRM* yielded increases in prediction accuracy that appeared moderately significant (hereafter “moderately significant” refers to *p* ≤ 0.05 based on the liberal “naïve” t*-*test) for PH (WS4U-C2, U4X-N) and St (Liberty-C2) (Table 4, Table S1). However, prediction accuracy for St (WS4U-C2) was lower, with a moderately significant difference. More intriguing is the consistent decrease in prediction accuracy with L4X-NE, with differences being small yet highly significant for PH and HD (hereafter “highly significant” refers to *p* ≤ 0.05 based on the conservative t*-*test adapted from Nadeau and Bengio 2003; see *Material and methods* for details), and moderately significant for St. It is unclear whether these differences are due to the consistently higher accuracies achieved with L4X-NE (in *Target*) compared to other populations, or a result of L4X-NE being relatively under-connected to the other populations in the sample (Figure S2, Figure 3). Both factors could very well contribute to the observed decreases in accuracy when incorporating information from the whole sample. Compared to *SPM-GRM, SPM-GLASSO* did not yield any notable increase in prediction accuracy, with generally similar accuracies, and differences from *SPM-GRM* ranging from −0.036 (for St, Liberty-C2) to +0.016 (for HD, WS4U-C2).

**Table 4.**
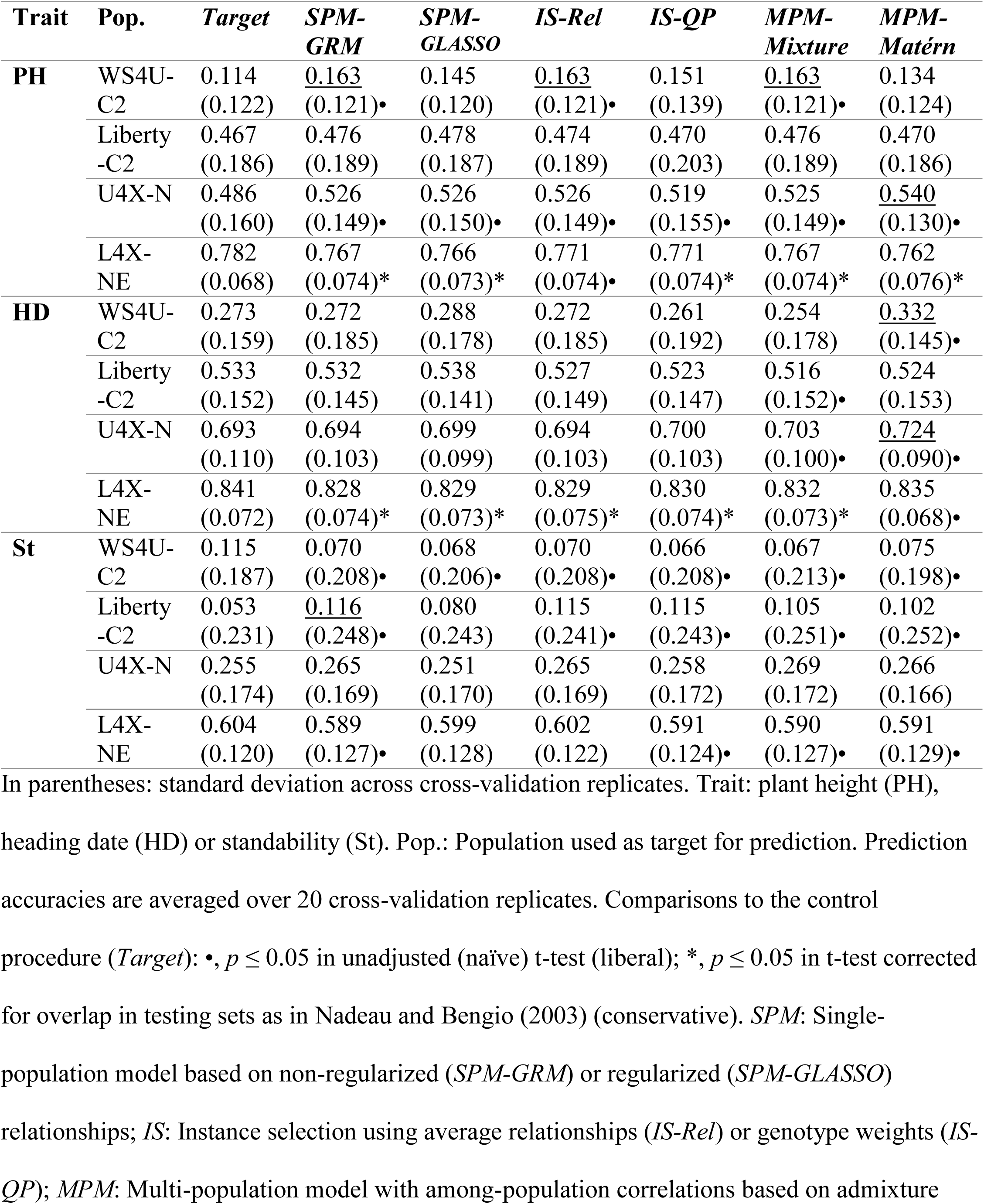
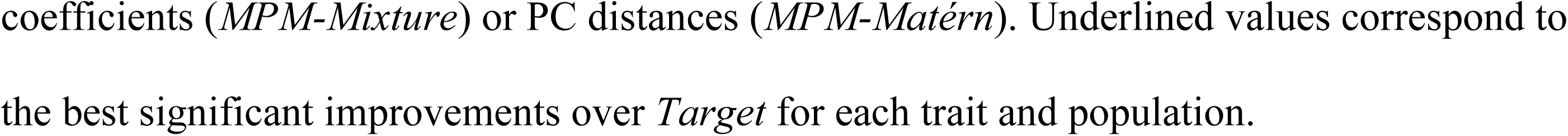
Average prediction accuracy by trait, target population and prediction procedure

### Instance selection in contexts of free and fixed CS size

In the fixed CS size context, increasing CS size often resulted in higher accuracy, with a plateau reached around the *Target* CS size, i.e., the number of individuals belonging to the sample population as the TS, which corresponded to 11%-15% of the available individuals (Figure 4; Table S2). The observed plateaus suggest that adding individuals from extraneous populations, without explicitly modelling heterogeneity, did not add useful signal for predictive ability of GBLUP; they also suggest that GBLUP is quite robust to such “superfluous” signal in a multi-population context. Exceptions were PH (WS4U-C2, U4X-N) and St (Liberty-C2) for which higher CS sizes did result in significant increases in accuracy compared to *Target*, with one highly significant increase achieved by *IS-Rel* with 75% selected individuals (Figure 4). Conversely, with L4X-NE for all traits and WS4U-C2 for St, adding more individuals actually deteriorated accuracy (Figure 4). The results about L4X-NE make sense in light of the facts previously noted for *SPM*: higher accuracies in L4X-NE and lack of kinship with other populations (Figure 3). When the CS was selected based on *IS-Rel*, CS sizes lower than the *Target* CS size resulted in significantly lower prediction accuracies compared to *Target*, except with WS4U-C2 (PH, St). Furthermore, at these low CS sizes, *IS-Rel* did not perform significantly better than random selection in some cases (Figure 5): PH (U4X-N, Liberty-C2), HD (U4X-N), and St (U4X-N, WS4U-C2). It was even significantly worse than random selection for St (Liberty-C2) (Figure 5). In comparison, *IS-QP* was often significantly better than *IS-Rel* at these low CS sizes, with substantial differences observed with U4X-N and L4X-NE (Figure S3). This advantage of *IS-QP* translated into similar accuracies compared to *Target* with 10% selected individuals, and small, sometimes non-significant, decreases with 5% selected individuals (Figure 4). Accordingly, *IS-QP* maintained its advantage over random selection at low CS sizes, with a consistent relative improvement as CS size decreased (Figure 5). One exception was St (Liberty-C2), for which there was a (non-significant) decrease in accuracy relatively to random selection with 5% selected individuals. At intermediate CS sizes (35%-85% of selected individuals), *IS-QP* performed similarly to *IS-Rel*, with a small (yet significant) advantage over *IS-Rel* for HD (L4X-NE, U4X-N) but a significant disadvantage in three cases (PH, WS4U-C2; HD, WS4U-C2; St, Liberty-C2) (Figure S3).

**Figure 4.**
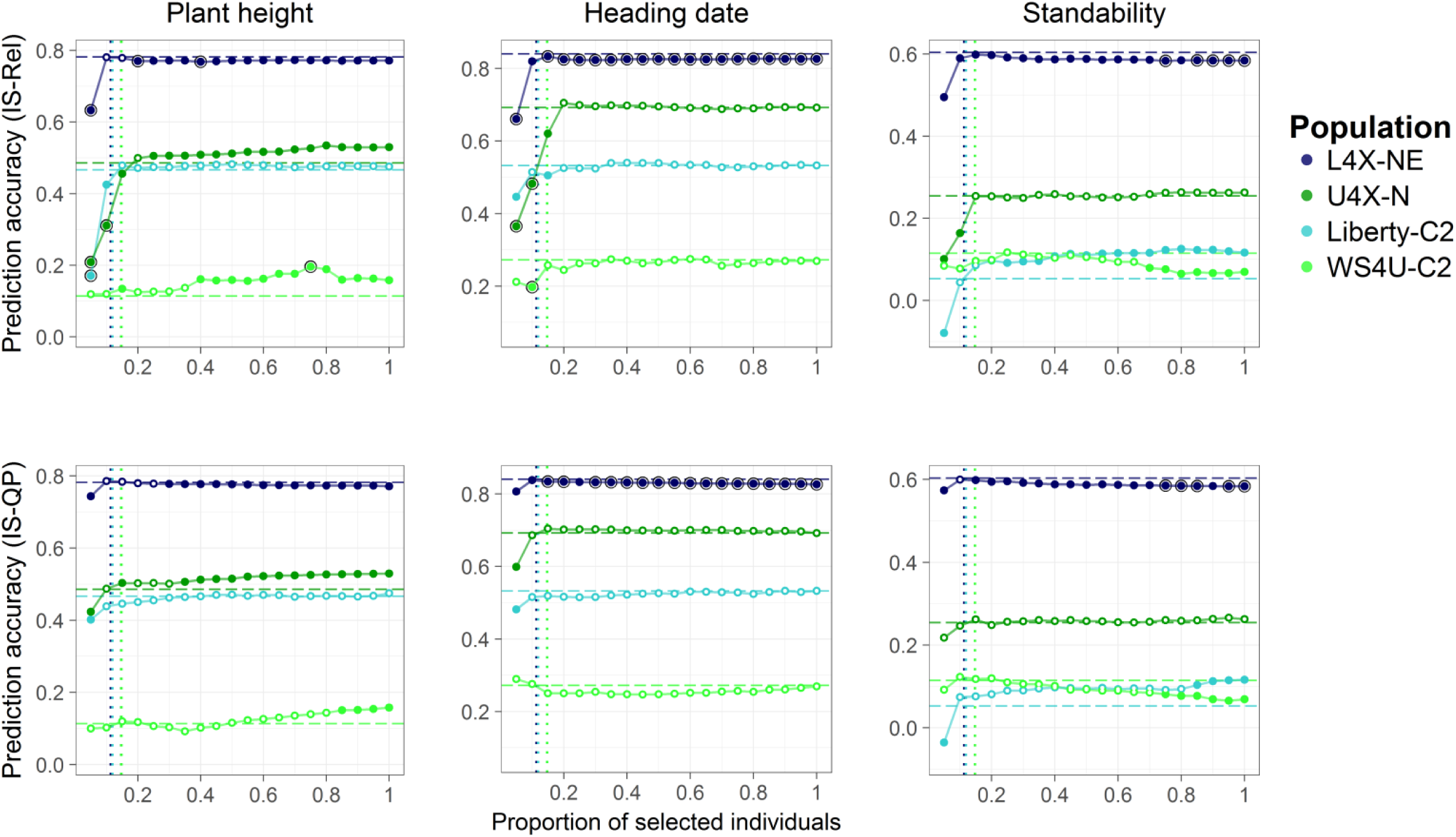
Average prediction accuracies from instance selection with fixed calibration set sizes Prediction accuracies (over cross-validation replicates) from instance selection are compared to the control procedure (*Target*). Upper panel: prediction accuracies from *IS-Rel*. Lower panel: prediction accuracies from *IS-QP*. Horizontal dashed lines indicate the average prediction accuracy from *Target*; vertical dotted lines indicate the corresponding proportion of individuals in the CS: 11%, 15%, 12% and 15% for L4X-NE, U4X-N, Liberty-C2 and WS4U-C2, respectively. Comparisons to the control: Colored fill, *p* ≤ 0.05 in unadjusted (naïve) t-test (liberal); black circle, *p* ≤ 0.05 in t-test corrected for overlap in testing sets as in Nadeau and Bengio (2003) (conservative).

**Figure 5.**
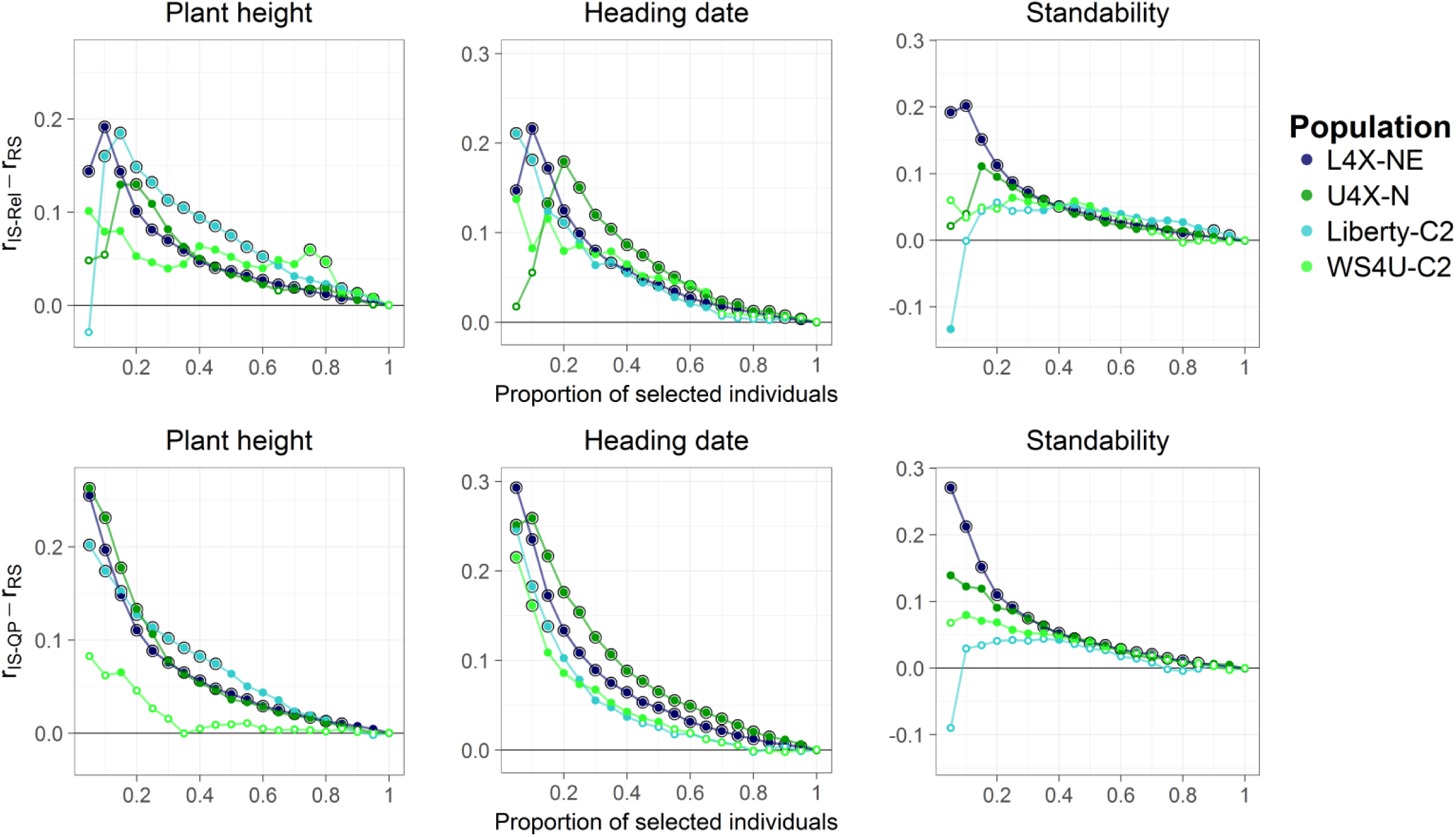
Differences in prediction accuracy between instance selection and random selection, with fixed calibration set sizes Upper panel: difference in mean prediction accuracy (over cross-validation replicates) between *IS-Rel* and random selection (*r*_*IS*–*Rel*_ – *r*_*RS*_). Lower panel: difference in mean prediction accuracy between *IS-QP* and random selection (*r*_*IS*–*QP*_ – *r*_*RS*_). Significance of differences: colored fill, *p* ≤ 0.05 in unadjusted (naïve) t-test (liberal); black circle, *p* ≤ 0.05 in t-test corrected for overlap in testing sets as in Nadeau and Bengio (2003) (conservative).

In a context of free CS size, *IS* procedures yielded some significant improvements over *Target*, similarly to *SPM-GRM*. In fact, both *IS-Rel* and *IS-QP* tended to select many, if not all, of the available individuals (Table 5). *IS-QP* tended to select less individuals than *IS-Rel*, except for HD (Liberty-C2, L4X-NE) and St (Liberty-C2, L4X-NE). However, the differences in selection behavior between *IS-Rel* and *IS-QP* generally mattered little, with the exceptions of PH (WS4U-C2), HD (WS4U-C2) and St (L4X-NE), for which *IS-QP* performed slightly worse than *IS-Rel* (−0.012, −0.011 and −0.011 in average accuracy, respectively). In the cases where *SPM* yielded significantly lower accuracies than *Target* (St with WS4U-C2, and all traits with L4X-NE), *IS* procedures failed to select an appropriately low number of individuals that would have prevented these decreases in accuracy (Figure 4), with the notable exception of *IS-Rel* for St (L4X-NE) (Table 4, Table 5).

**Table 5.**
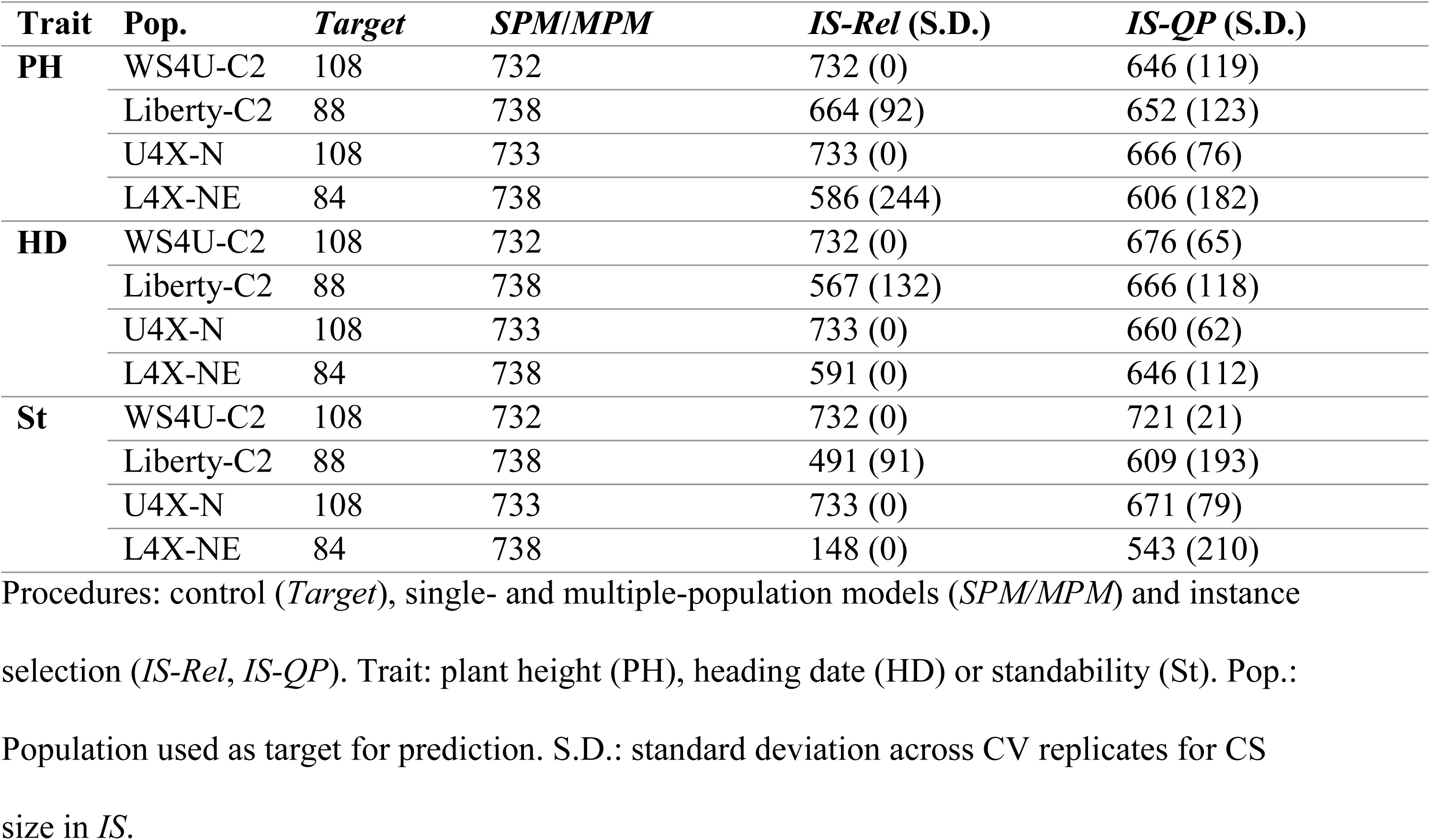
Size of the calibration set in the different procedures

| **Trait** | **Pop.** | **Target** | **SPM/MPM** | **IS-Rel (S.D.)** | **IS-QP (S.D.)** |
| --- | --- | --- | --- | --- | --- |
| **PH** | WS4U-C2 | 108 | 732 | 732 (0) | 646 (119) |
|  | Liberty-C2 | 88 | 738 | 664 (92) | 652 (123) |
|  | U4X-N | 108 | 733 | 733 (0) | 666 (76) |
|  | L4X-NE | 84 | 738 | 586 (244) | 606 (182) |
| **HD** | WS4U-C2 | 108 | 732 | 732 (0) | 676 (65) |
|  | Liberty-C2 | 88 | 738 | 567 (132) | 666 (118) |
|  | U4X-N | 108 | 733 | 733 (0) | 660 (62) |
|  | L4X-NE | 84 | 738 | 591 (0) | 646 (112) |
| **St** | WS4U-C2 | 108 | 732 | 732 (0) | 721 (21) |
|  | Liberty-C2 | 88 | 738 | 491 (91) | 609 (193) |
|  | U4X-N | 108 | 733 | 733 (0) | 671 (79) |
|  | L4X-NE | 84 | 738 | 148 (0) | 543 (210) |

Procedures: control (*Target*), single- and multiple-population models (*SPM/MPM*) and instance
selection (*IS-Rel*, *IS-QP*). Trait: plant height (PH), heading date (HD) or standability (St). Pop.:
Population used as target for prediction. S.D.: standard deviation across CV replicates for CS
size in *IS*.

### Multi-population models and marker-by-population interactions

The inferred mixing parameter *ρ* from the *MPM-Mixture* model was null (or close to null), low and intermediate, for PH, St and HD respectively, with estimations being quite consistent over CV replicates (Table 3). Expectedly, the improvement in fit, relatively to *SPM-GRM*, was non-significant for PH, rather significant (*p* < 0.05) for St, and strongly significant (*p* < 0.001) for HD (Table 3). In *MPM-Matérn*, the inferred correlation functions differed substantially across traits, while being quite consistent over CV replicates (Table 3, Figure 6): *k*_*v,h*_ roughly resembled an exponential kernel with PH and HD, and was more similar to a Gaussian kernel with St, for which a “shoulder” maintained high correlation in marker effects for individuals that were relatively close to each other, based on their PCs. Inferences regarding **Ω**_*n*_ in *MPM-Matérn* were weakly significant for PH and St, with *p*-values sometimes above 0.05 (ranging from 0.007 to 0.100 for PH and from 0.014 to 0.062 for St); in contrast, inferences regarding **Ω**_*n*_ for HD were strongly significant (Table 3). Interestingly, distances based on PCs may be equivalent to distances based on allele frequencies. Specifically, 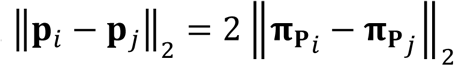, where 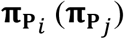 is the *m*-vector of individual-specific allele frequencies of individual *i* (*j*) as described by Conomos *et al.* (2016), with population structure described by [**1**_*n*_ **P**] (Appendix A3). Therefore, the significant relationship between PC-based distances and correlations in marker effects (depicted by **Ω**_*n*_) for HD in *MPM-Matérn* indicates that marker effects for this trait were highly sensitive to variation in allele frequencies across genetic backgrounds.

**Figure 6.**
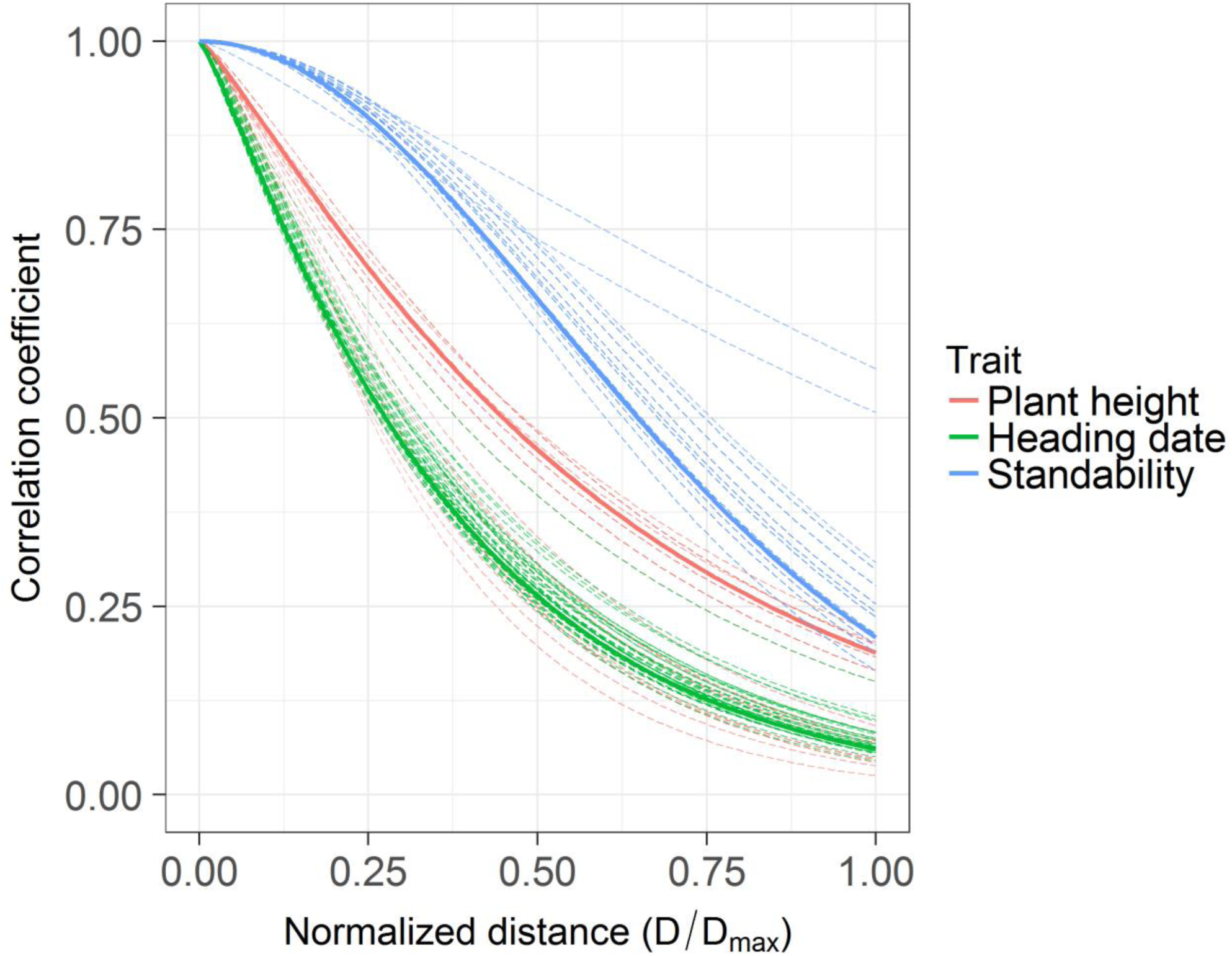
Shape of the inferred correlation functions in *MPM-Matérn* Solid curves: inferred correlation function in the whole sample; dashed curves: inferred correlation functions in cross-validation replicates. Correlations are functions of *D* = ‖**p**_i_ – **p**_j_‖_2_(the Euclidean distance between population-structure PCs for any pair of individual (*i,j*)), scaled by *D*_*max*_, the maximum distance ‖**p**_i_ – **p**_j_‖_2_ observed over pairs.

Regarding prediction accuracy, the performance of *MPM-Mixture* was very similar to that of *SPM-GRM*, with differences in accuracy ranging from −0.018 to +0.009 (Table 4). Quite surprisingly, *MPM-Mixture* displayed slightly deteriorated accuracies for HD (with the exception of U4X-N), despite the strongly significant improvement in fit for this trait. In contrast, *MPM-Matérn* yielded larger differences in accuracy, ranging from −0.019 to +0.060 (Table 4). With the two upland target populations (WS4U-C2 and U4X-N), noteworthy increases in prediction accuracy (+0.060 and +0.030 respectively) were observed for HD. In these two cases, moderately significant differences in accuracy compared to *Target* could be achieved, while no significant improvement could be obtained from *SPM-GRM*. With the two other target populations (Liberty-C2 and L4X-NE), smaller differences in accuracy (−0.008 and +0.007 respectively) were observed for HD. Our results suggest that a very high increase in quality of fit, as was observed for HD with *MPM-Matérn*, may allow for an increase in accuracy, but with no absolute guarantee. In the analysis of Heslot and Jannink (2015) across various multi-population contexts, there seemed to be a positive relationship between differences in quality of fit, as measured by the Akaike information criterion (AIC), and differences in prediction accuracy. Although this relationship was quite loose, it could be noted that for very high increases in AIC (≥ 30), gains in accuracy were null to high, similarly to the situation of *MPM-Matérn* with HD, for which increases in AIC varied from 28.68 to 42.88 across CV replicates (Table 3). Therefore, stringent thresholds on AIC increases could probably be used in *MPM* to avoid relative decreases in accuracy. Other characteristics and guidelines, related to the sample or the fitted model, may also be useful for this type of indication.

## DISCUSSION

### Conclusions and comments

The present study assessed various procedures to accommodate population heterogeneity in diverse samples, with an application in switchgrass. We employed three typical strategies for dealing with marker-by-population interactions, i.e., ignoring (*SPM*), reducing (*IS*), or modelling (*MPM*) the source of heterogeneity in the data. These general strategies had previously been mentioned, e.g. by Bernardo (2002) about the analysis of genotype-by-environment interactions.

Here *SPM* often seemed robust to population heterogeneity, regarding prediction accuracy (Table 4). This robustness was certainly due to the ability of GBLUP models to combine information from individuals according to the specified covariance matrix, by transferring information preferentially from the more related individuals (Searle *et al.* 2006, Habier *et al.* 2013). Furthermore, GBLUP models were probably all the more robust that marker density was high, which translated into high estimation accuracy of genomic relationships (Casella and Berger 2002, Endelman and Jannink 2012). However, some decreases in accuracy compared to the control procedure (*Target*) suggest that robustness of *SPM* may have been affected by other factors. Such factors may be related to relationships within the sample, e.g. under-connectedness of some populations with others (Figure 3), or differences in accuracy of the prediction model from one population to another, in a single-population context (Table 4). Our proposed procedure (*SPM-GLASSO*), relying on a regularized form of the genomic relationship matrix, did not yield any improvement in prediction accuracy compared to the standard procedure (*SPM-GRM*). However, *SPM-GLASSO* was useful for inferring graphs of relationships within the sample, conditionally on population structure, which were used to derive informative features about our sample (Figure 3, Figure S2).

Selecting individuals in *IS* was not useful for improving prediction accuracy compared to *SPM*, in a context of free CS sizes (i.e., when *IS* is used as means of optimization with respect to the CS). However, *IS* procedures were useful compared to random selection in contexts of fixed CS sizes, i.e., when restrictive numbers of individuals were used for calibration (Rincent *et al.* 2012, Isidro *et al.* 2014, Akdemir *et al.* 2015). In this type of scenarios, with small CS sizes (less individuals than in *Target*), the proposed procedure (*IS-QP*), which not only accounted for relationships between the CS and the TS but also redundancy within the CS, was particularly useful (Figure 5). In comparison, the more typical approach (*IS-Rel*), which only accounted for relationships to the TS, tended to lose its advantage over random selection as CS size decreased (Figure 5). The relative superiority of *IS-QP* at low CS sizes is consistent with the findings of Pszczola *et al.* (2012), which suggested that redundancy within the CS was detrimental to prediction accuracy, for a given level of relationships to the TS.

In our case study, *MPM* procedures yielded highly significant improvements in fit for one of the three traits assayed (HD), in comparison to *SPM* (Table 3). Our proposed procedure (*MPM-Matérn*) relied on non-linear kernel functions for estimating population-level correlations in **Ω**_*n*_, and was the only procedure to be more accurate than *SPM* for HD (Table 4). Differences in accuracy from *SPM* to *MPM* were smaller and seemed less predictable for PH and St, as could be expected from the more modest improvements in fit for these two traits. Our results exemplify the potential usefulness of parsimonious multi-population models, which are all the more interesting that they can be applied on samples comprising many populations. In contrast, typical multi-trait models would be computationally intractable or statistically inefficient here, since those would rely on one parameter for each population pair to model correlations among populations in **Ω**_*n*_ (e.g., 21 parameters for *K* = 7 population clusters).

In this study, marker data was based on exome capture sequencing, which targets a selected subset of exons for sequencing and subsequent SNP calling (Hirsch *et al.* 2014, Evans *et al.* 2014). The potential lack of representation of causal variants by our assay may have resulted in loss of prediction accuracy. While total lack of representation of some regions imposes a limit on prediction accuracy achievable by our procedures, the relative overrepresentation of some genomic regions could be, to some extent, alleviated by genomic relationships which account for correlation among markers and differential degree of tagging of loci in the marker data (Speed *et al.* 2012, Ramstein *et al.* 2016, Wang *et al.* 2017).

### Potential of regularization in genomic analyses

In our study, the lack of benefit from regularization was probably due to the high marker density (Endelman and Jannink 2012, Müller *et al.* 2015). In studies with lower marker densities, in which genomic relationship estimates are less accurate, *SPM-GLASSO* may have been more useful compared to *SPM-GRM*. Specifically, based on normal theory on sample covariance matrices, when the number of markers is of the same order of magnitude or lower than the number of individuals (*m* ≤ *n*), the estimated genomic relationships matrix **G** is expected to be inaccurate (in the sense that its eigenvalues are estimated with large errors) and therefore may benefit from regularization (Johnstone 2001, Bickel and Levina 2008, Fan *et al.* 2013). So regularization would be particularly useful in applications with few markers (e.g., Lorenz and Smith 2015: *n* = 1068, *m* = 342 in barley) or many individuals (e.g., Lourenco *et al.* 2015: *n* = 51,883, *m* = 38,528 in beef cattle). Further analyses assessing the performance of our proposed regularization in these contexts would be useful. In such studies, *SPM-GLASSO* would need to be compared to other regularization techniques previously proposed (Endelman and Jannink 2012, Müller *et al.* 2015). Such techniques were generally based on shrinkage to a target matrix **T**. In presence of population structure, we recommend having **T** depict relationships at the population level (e.g., **T** = **BB**^′^, rather than **T** = **I**) and shrinking relationships conditioned on population structure (e.g., **G**_*B*_ = **G** – **BB**^′^). Only then would relationships under regularization conform to the assumption of (conditional) sparsity, i.e., most relationships assumed to be zero, which is a critical assumption for proper and meaningful regularization of covariance matrices (Figure 2; Fan *et al.* 2013). Regularization of relationships may also hold promise for other applications than genomic prediction. In particular, regularization may be useful in kinship inference, e.g., in parentage testing (Wang 2012) and control of inbreeding, in species conservation (Allendorf *et al.* 2010) or animal breeding (Pryce *et al.* 2012). In this context, kinships under regularization should probably be those proposed by Conomos *et al.* (2016), similar to the conditional relationships defined in this study, but scaled individually, so as to better estimate kinship coefficients conditionally on population structure. In kinship inference, regularization by the graphical LASSO would be useful, not only to increase estimation accuracy, but also to infer pedigrees. In this type of studies, choice of *λ* would be based, not on REML in a prediction model, but on selection techniques employed on the covariance matrix, e.g. based on cross-validation (Bickel and Levina 2008), information criteria (Foygel and Drton 2010), or stability of inference (Liu *et al.* 2010). The inferred pedigrees may then be used to depict kinships and make decisions accordingly, but may also be used in pedigree-based linkage analyses (Day-Williams *et al.* 2011), or as input to other procedures (e.g. in *IS*, as mentioned below).

### Improvement of procedures

The regularization method applied here in *SPM-GLASSO* imposed sparsity on the inverse matrix of conditional genomic relationships, thereby inferring a graph of recent relationships among individuals in the panel. Other regularization techniques act directly on the covariance matrix. Those include various thresholding methods (Rothman *et al.* 2009, Cai and Liu 2011), which may be useful, especially when relationships are derived from low-density markers. However, such methods do not necessarily guarantee positive definiteness of the regularized relationship matrices, which could be an issue when using them in linear mixed models. One other aspect of regularization that could be improved is the selection of the regularization parameter (*λ*). Here, we chose to select *λ* based on REML for a given outcome, but other selection techniques employed on the covariance matrix (as mentioned above) may be more relevant. Further research would be necessary to explore the potential of such selection techniques for improving regularization of genomic relationship matrices with respect to prediction accuracy.

In a context of unrestricted CS sizes, the tendency of *IS* procedures to select too many individuals, even when this was detrimental to prediction accuracy, may have been due to an overestimation of accuracy by CD_mean_ with larger CS sizes (Table 4, Table 5, Figure 4). Based on the results of Hayes *et al.* (2009c), this upward bias in CD_mean_ may be of particular concern in a multi-population context. Therefore, other metrics than CD_mean_ may result in more pertinent selections in *IS*. Another improvement of *IS* may come from selection of individuals for prediction of each TS individual considered separately, as was proposed by Lorenz and Smith (2015). Because the *IS* procedures would then be run for one TS individual at a time, computationally intensive procedures based on stochastic algorithms would certainly not be applicable, making the *IS* procedures presented here all the more relevant. Finally, *IS* could be further improved by using other types of relationships than those used here. For example, as was recommended by Pszczola *et al.* (2012) and Wientjes *et al.* (2013), selections could be based on squared relationships, i.e., **G** ○ **G** instead of **G** in *IS-Rel* (and by analogy, **XX**^′^○ **XX**^′^instead of **XX**^′^in *IS-QP*). Alternatively, entries in the relationship matrix could be replaced with those inferred in *MPM*, e.g. using **Ω**_*n*_ ○ **XX**^′^instead of **XX**^′^in *IS-QP*. Population heterogeneity would then be accounted for when selecting individuals from different genetic backgrounds. Finally, *IS* could rely on graphs of relationships such as those inferred from *SPM-GLASSO*. Selection of individuals would then be based on measures of connectivity between available individuals and the TS. Such measures could be the lengths of average shortest paths between each individual and the TS, or graph-based kernel functions, e.g. derived from the number of edges connecting each individual and the TS (Bishop 2006).

Multi-population models were generally not useful when the improvement in model fit was modest. Therefore, a possible improvement of *MPM* procedures could simply come from model selection as an integral part of the fitting process, based for example on the Bayesian information criterion (BIC). In fact, the BIC differences relative to *SPM-GRM* were almost always negative for PH and St in *MPM* (Table 3). For these two traits, differences in prediction accuracy from *SPM* to *MPM* were quite inconsistent, especially with *MPM-Matérn*, so model selection could probably have made *MPM* procedures more robust. Another way of potentially improving *MPM* procedures would be to use other types of kernels than those used here. For example, one may use linear kernels based on population-level covariates (e.g. PCs) in place of **AA**^′^in *MPM-Mixture*, hence taking an approach similar to that of Jarquín *et al.* (2014) who modelled genotype-by-environment interactions through environmental covariates, in multi-environment genomic prediction models. Besides, modelling resemblance among population clusters in *MPM-Mixture*, by **AΘ**_*K*_**A**^′^in place of **AA**^′^(where **Θ**_*K*_ captures similarity based on metrics at the population level), could be useful to increase quality of fit, and possibly prediction accuracy. Also, the relationship matrix used in *MPM* (**Ω**_*n*_ ○ **XX**^′^) may be regularized, thereby shrinking further – or even setting to zero – the relationships that had already been reduced through **Ω**_*n*_. Finally, an interesting way of extending the *MPM* procedures described here would be to incorporate more information at the population level. Here in *MPM*, population homogeneity was captured through admixture coefficients (*MPM-Mixture*) or differences in PC coordinates (*MPM-Matérn*), the latter reflecting differences in allele frequencies (Appendix A3). However, marker-by-population interactions may also be due to differences in LD patterns (Wientjes *et al.* 2016). Therefore metrics depicting such differences could be particularly appropriate for capturing population heterogeneity. Further research would be necessary to determine the type of metrics to use for reflecting differences in LD patterns, and the appropriate way to parsimoniously combine the different types of information regarding population differentiation in *MPM*. Interestingly, geographical distance may succinctly depict population differentiation, due to differences in allele frequencies and/or differences in LD patterns. Fitting population-level correlations **Ω**_*n*_ as a function of distance of origin would then be particularly useful in species under strong geographical structure, which include switchgrass (Grabowski *et al.* 2014), but also human (Coop *et al.* 2009), as was clearly shown in samples from Europe (Novembre *et al.* 2008), Africa (Bryc *et al.* 2010) and Latin America (Ruiz-Linares *et al.* 2014). Models such as *MPM-Matérn*, which are parsimonious yet flexible in the shape of the fitted correlation function (Figure 6), are promising in various applications on diverse samples, in prediction studies, but also in inferential studies aiming at characterizing the basis for population differentiation.

### Applications and prospects

Based on our case study, we would recommend using *MPM* whenever a strong improvement in model fit is achieved. Otherwise *SPM* would be the method of choice, since it is often robust enough to perform at least as well as *Target*. However, *Target* may be preferred when making predictions on “outlier populations” such as L4X-NE, which are under-connected to other populations and are characterized by relatively higher prediction accuracy in a single-population context. Only when the CS sizes are restricted (fixed) would *IS* procedures be useful – even though further improvements may make *IS* more competitive in contexts of free CS sizes. In such situations, we recommend using *IS-QP* instead of *IS-Rel*, especially when the CS size ought to be small. Nevertheless, more empirical studies on population heterogeneity would have to follow to support the conclusions from our specific application. Such studies could apply to various contexts: in particular, predictions on diverse samples and dynamic breeding programs. The former includes analyses similar to our case study as well as analyses on more complex data, such as historical datasets, in which not only population heterogeneity but also genotype-by-environment interactions must be taken into account (Dawson *et al.* 2013, Rutkoski *et al.* 2015). The latter involves selection across multiple breeding generations, which might not necessarily suffer from strong population heterogeneity (Sallam *et al.* 2015, Auinger *et al.* 2016) but could nonetheless benefit from robust multi-population models for potential increase in persistency of accuracy over generations (Habier *et al.* 2007). In this context, *IS* procedures could also be interesting, for example if a subset of non-selected individuals may be assayed phenotypically during the breeding program. In the context of diverse samples or dynamic breeding, simulation studies could also be useful for assessing the suitability of procedures to accommodate population heterogeneity. Differences in allele frequencies and differential LD patterns could be simulated by various genealogies, as was done for example by De Roos *et al.* (2009). Additionally, dependency of marker effects on allele frequencies could be simulated, not only via allele fixation in specific populations, but also by underlying non-additive genetic effects. Indeed, additive marker effects, as captured by linear models such as GBLUP (which typically do not account for interactive marker effects), depend on allele frequencies at the loci with which they interact (Hill *et al.* 2008, Mäki-Tanila and Hill 2014, Hill and Mäki-Tanila 2015). Therefore, dominance and epistatic effects could be simulated to generate dependency of additive marker effects on allele frequencies, which may then be captured by methods such as *MPM-Matérn* (Appendix A3). Though investigations based on simulations would be complex and, to some extent, arbitrary by their choice of genealogies and genetic architectures, they would provide useful frameworks for assessing procedures, such as those presented in this study, in contexts of population heterogeneity.

## ACKNOWLEDGEMENTS

We are grateful to Jeremy Schmutz of the Department of Energy Joint Genome Institute and Hudson Alpha for his work on the switchgrass genome. We are also grateful to Nick Baker and Joseph Halinar, USDA-ARS, Madison, WI for assistance with field operations and data collection. Finally, we would like to thank two anonymous reviewers for their constructive comments. This research was funded in part by the following agencies and organizations: the U.S. Department of Energy Great Lakes Bioenergy Research Center, DOE Office of Science BER DE-FC02-07ER64494 (laboratory operations, genotyping, and bioinformatics), the U.S. Department of Energy Joint Genome Institute supported by the Office of Science of the U.S. Department of Energy under Contract No. DE-AC02-05CH11231 (sequencing), Agriculture and Food Research Initiative Competitive Grant No. 2011-68005-30411 from the USDA National Institute of Food and Agriculture (CenUSA; field operations and phenotypic measurements), USDA-ARS Congressionally allocated funds (field operations, technical support, and logistics), and the University of Wisconsin Agricultural Research Stations (field operations). Mention of commercial products and organizations in this manuscript is solely to provide specific information. The USDA is an equal opportunity provider and employer. The funders had no role in study design, data collection and analysis, decision to publish, or preparation of the manuscript.

### Appendix A1 – Equivalence of fit in linear mixed models after regressing out fixed-effect variables from random-effect variables

In this section, matrix notations are not consistent with those in the main text; **I** refers to the identity matrix with dimensions equal to the number of observations.

Consider the following two models:

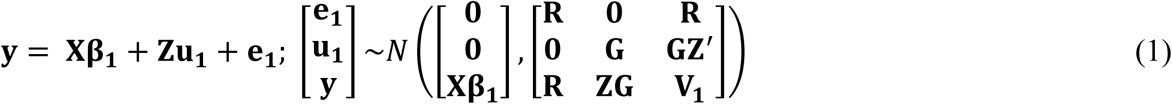

where **V**_1_ = **ZGZ**^′^+ **R**

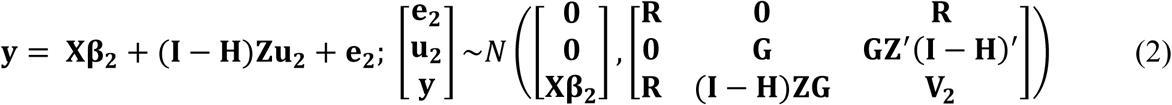

with **H** = **X**(**X**′**R**^**–1**^**X**)^**–1**^**X**′**R**^**–1**^ and **V**_**2**_ = (**I** – **H**)**ZGZ**^′^(**I** – **H**)^′^+ ***R***.

For given **R** and **G** (possibly estimated by ML or REML), in model (1), the ML estimates of regression coefficients are 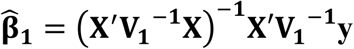 (as best linear unbiased estimators, BLUEs) and 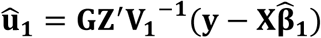 (as best linear unbiased predictors, BLUPs); in model (2), the mixed model equations (MME) are

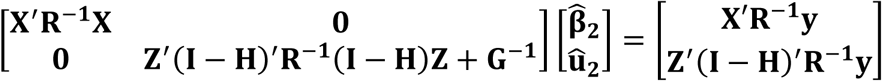

so the ML estimates of regression coefficients are (as solutions of the MME) 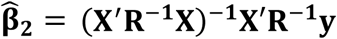 and **û**_2_ = (**Z**^′^(**I** – **H**)^′^**R**^**–1**^(**I** – **H**)**Z** + **G**^**–1**^)^**–1**^**Z**^′^(**I** – **H**)^′^**R**^−1^**y** (Henderson, 1984).

□ For given **R** and **G**, fitting model (1) and fitting model (2) by ML are equivalent, in that **ê**_1_ = **ê**_2_, so **ŷ**_1_ = **ŷ**_2_.

Consider the two matrices **P** and **S** such that **P** = **V1**^−1^ – **V1^−1^X**(**X**′**V1^−1^X**)**^−1^X**′**V1^−1^** and **S** = **R**^**–1**^ – **R**^**–1**^**X**(**X**^′^**R**^**–1**^**X**)^**–1**^**X**′**R**^**–1**^.

As shown by Searle *et al.* (2006, pp. 282-283):

- **P = S − SZ(Z′SZ + G^−1^)^−1^Z′S**

Therefore:

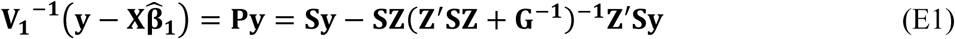

Moreover:

- 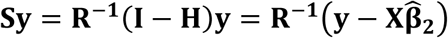
- Because **S** = **R**^**–1**^(**I** – **H**) = (**I** – **H**)^′^**R**^**–1**^= (**I** – **H**)^′^***R***^**–1**^(**I** – **H**), **SZ** = **R** ^−1^ (**I – H**)**Z** and **Z** ′ **SZ** = **Z** ′ (**I – H**) ′ **R** ^−1^ (**I – H**)**Z**.

Therefore (E1) simplifies into 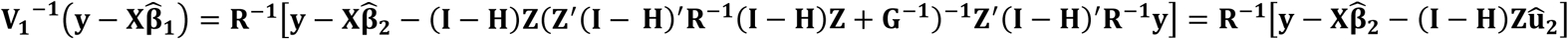. So:

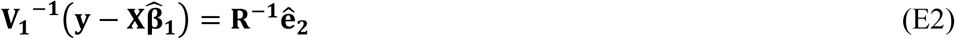

Besides, as shown by Henderson (1984, Chapter 5 p. 9) by application of the Sherman-Morrison-Woodbury formula to a general variance formulation:

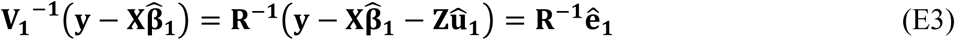

Since **R**^−1^ is positive definite (so it is full rank), it follows from (E2) and (E3) that **ê**_1_ = **ê**_2_.

▪

□ For given **R** and **G**, **û**_1_ = **û**_2_.

As previously stated:

- **S = (I − H)′R^−1^(I − H)**

Therefore, it follows from (E1) that:

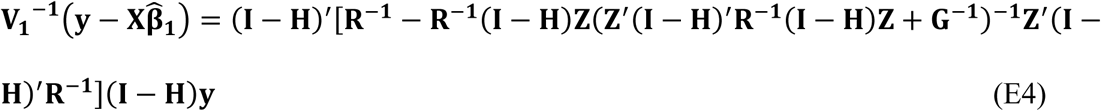

By the Sherman-Morrison-Woodbury formula, **V_2_^−1^** = [**R** + (**I** – **H**)**ZGZ**^′^(**I** – **H**)^′^]^**–1**^ = [**R**^**–1**^ – *R*^−1^(**I** – **H**)**Z**(**Z**^′^(**I** – **H**)^′^**R**^**–1**^(**I** – **H**)**Z** + **G**^**–1**^)^**–1**^**Z**^′^(**I** – **H**)^′^**R**^**–1**^].

So it follows from (E4) that:

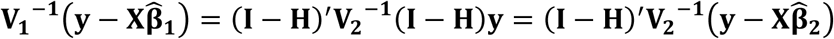

Therefore, as BLUP, **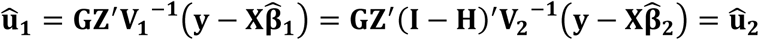**. ▪

### Appendix A2 – Multi-population GBLUP models for heterogeneous calibration sets

In this section, **1**_*t*_, **I**_*t*_, and **J**_*t*_ refer to the vector of ones, identity matrix, and matrix of ones, respectively, of dimensions *t, t* ×; *t* and *t* ×; *t* (where *t* is specified).

Consider the following model for population-specific marker effects with respect to *K* populations and *n* genotypes:

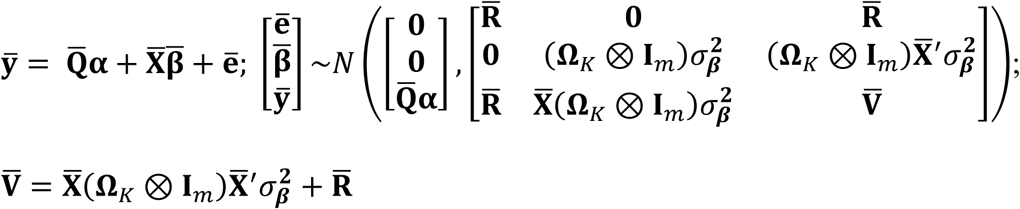

where ⊗ indicates the Kronecker product; 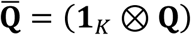 is the *Kn* x *p* design matrix for the *p*-vector**α** of fixed effects; 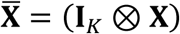 is the *Kn* x *Km* marker-data matrix for the *Km*-vector 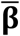 of marker effects at each of the *K* populations, with variance 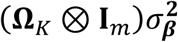. The matrix **Ω**_*K*_reflects covariances in marker effects between populations. The *Kn*-vector 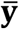, containing the phenotypic values for the *n* genotypes at each of the *K* populations, is hypothetical (and ill-defined from a practical standpoint), since genotypes typically do not belong to more than one population. The *Kn*-vector of residuals 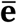, with unspecified variance 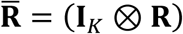, is assumed to be uncorrelated to marker effects 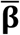.

Let 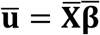 be the *Kn*-vector of additive genetic effects at each of the *K* populations,

As a linear combination of a normally-distributed vector 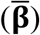, 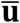 follows a normal distribution with expectation and variance as follows (Lehermeier *et al.*, 2015):

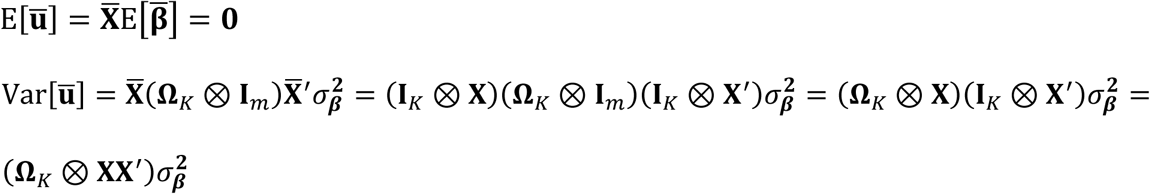

So a multi-population model for breeding values that is equivalent to the model described above, by identical mean and variance structures, is as follows:

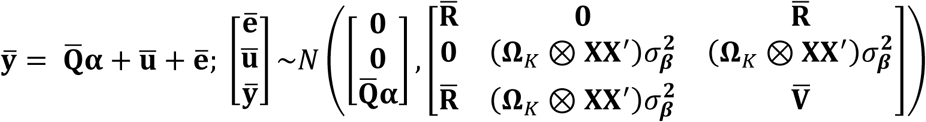

Now assume that *K* = *n*, and each population correspond to the specific genetic background of each individuals separately. By considering only observations at every individual’s specific genetic background, the above model reduces to:

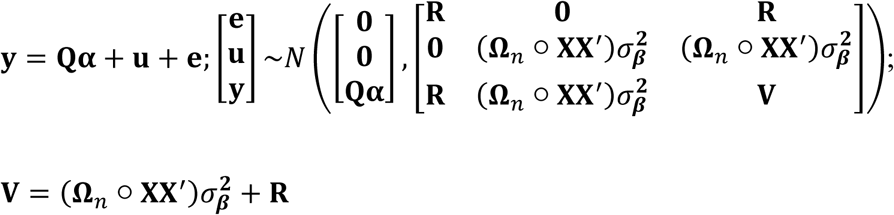

where ○ is the Hadamard product; **y** is the typical *n*-vector of observed phenotypic values; **u** and **e** are the corresponding additive genetic effects and residuals, respectively. Individual-specific marker effects are therefore accounted for by multiplying each element of the relationship matrix **XX**^′^ by the corresponding element of **Ω**_*n*_, thereby reflecting correlations in marker effects among individuals’ genetic backgrounds.

In general, we propose to infer **Ω**_*n*_ = (*ω*_*ij*_)_*n*×;*n*_ by *ω*_*ij*_ = *κ* (*φ*(**x**_*i*_), *φ*(**x**_*j*_)), where *φ* is some function of the *m*-vectors of marker variables x_*i*_ and x_*j*_, for any pair of individuals *i* and *j*, and *κ* is a valid kernel function guaranteeing that **Ω**_*n*_ be positive semi-definite. In *MPM-Mixture, φ*(**x**_*i*_) = **a**_*i*_ (*K*-vector of admixture coefficients for *i*) for any individual *i*, and the kernel function is 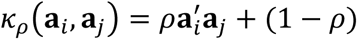, so that **Ω**_*n*_ is a “mixture” between a matrix of correlations restricted to population clusters and a matrix allowing full exchange of information across clusters, as in a standard GBLUP model. More generally, one could define the kernel function as 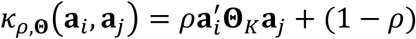, where **Θ**_*K*_ is a *K* x *K* matrix depicting relationships among clusters. Here, we simply set **Θ**_*K*_ = **I**_*K*_ and adjusted the kernel function (by REML) for *ρ* only.

In *MPM-Matérn, φ*(**x**_*i*_) =**p**_*i*_ (*d*-vector of PC coordinates for *i*) for any individual *i*, and the kernel function is a Matérn function *k*_*v,h*_(**p**_*i*_, **p**_*j*_) of ‖**p**_*i*_ – **p**_*j*_‖_2_, where ‖.‖_2_ is the Euclidean norm. Notably, it can be shown that ‖**p**_*i*_ – **p**_*j*_‖_2_ is proportional to 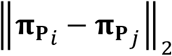, where 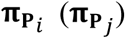 is the *m*-vector of individual-specific allele frequencies for individuals *i* (*j*), defined by projection of matrix **X** onto the column space of **Q**_**P**_ = [**1**_*n*_ **P**] (Appendix A3). So ‖**p**_*i*_ – **p**_*j*_‖_2_, which reflects differentiation with respect to coordinates at the leading PCs of **X**, also reflects differentiation with respect to individual-specific allele frequencies, with an underlying population structure represented by the same PCs. The allele frequencies 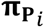 have been introduced by Conomos *et al.* (2016), in a study where they also recommended using principal components from a subset of unrelated individuals in **X**. Here, we simply applied PCA on the whole matrix **X**.

### Appendix A3 – Relationship between distance based on principal components and distance based on individual-specific allele frequencies

In this section, **1**_*t*_, **I**_*t*_, **J**_*t*_ and **0**_*s*×;*t*_ refer to the vector of ones, identity matrix, matrix of ones and matrix of zeros, respectively, of dimensions *t, t* ×; *t, t* ×; *t* and *s* ×; *t* (where *s* and *t* are specified).

We will consider the case where the PC matrix **P** consists of the first *d* PCs of **X**, and individual-specific allele frequencies are defined as (Conomos *et al.* 2016):

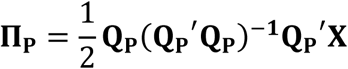

where **Q**_**P**_ = [**1**_*n*_ **P**] represents population structure through an intercept and the effects of the first *d* PCs of **X**. Vector 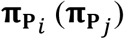 then consists of individual-specific allele frequencies (with respect to **Q**_**P**_) for individual *i* (*j*), such that:

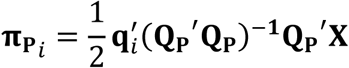

and similarly for 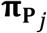 (**q**_*i*_ refers to the (*d* + 1)-vector of population-structure variables from **Q**_**P**_for individual *i*).

□ We will show that 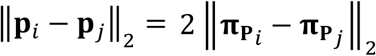 for any pair (*i, j*), i.e., Euclidean distances based on *d* PCs are equivalent, by proportionality, to those based on *m* individual-specific allele frequencies, with such frequencies as defined above.

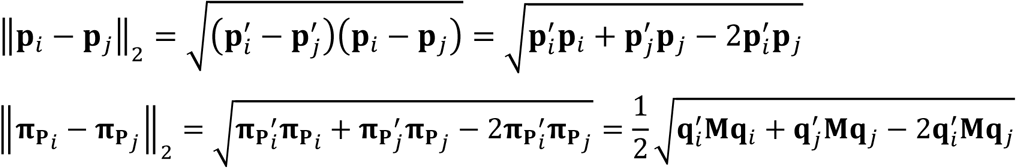

where M = (**Q**_**P**_^′^**Q**_**P**_)^−1^**Q**_**P**_^′^**XX**^′^**Q**_**P**_(**Q**_**P**_^′^**Q**_**P**_)^−1^.

Below, we will specify **M** more explicitly, to subsequently show that 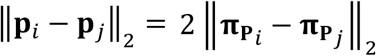.

Let 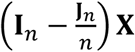 be the matrix of marker variables centered around their respective overall mean. Assuming *m* ≥ *n*, by eigenvalue decomposition 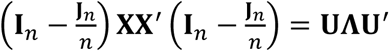, with **U** the *n* ×; *n* matrix of eigenvectors and Λ the *n* ×; *n* diagonal matrix of eigenvalues of 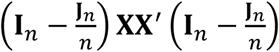 and 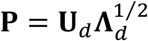, where the **U**_*d*_ is the *n* ×; *d* matrix of leading eigenvectors and 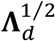 is the *d* ×; *d* diagonal matrix of corresponding singular values, assumed strictly positive.

Because **U**_*d*_ consist of left-eigenvectors of a column-centered matrix (associated with strictly positive eigenvalues),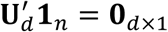 so 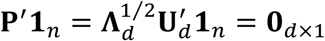.

Besides, **P**^′^**P** = **Λ**_*d*_.

So:

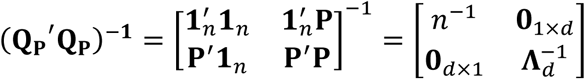

Moreover 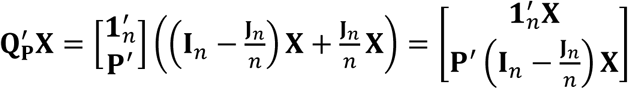 because 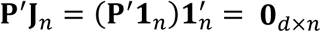.

So:

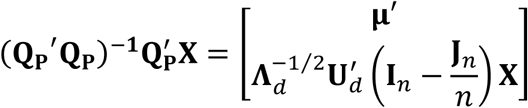

where **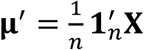**

Finally,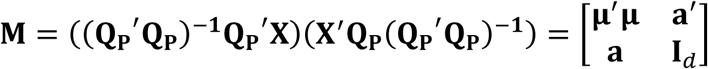
with:

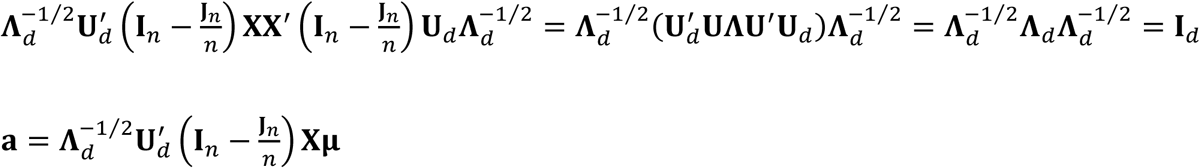

Therefore, for any pair of individuals (*i, j*):

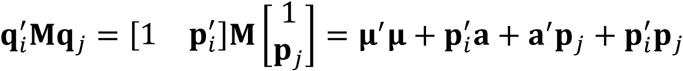

So:

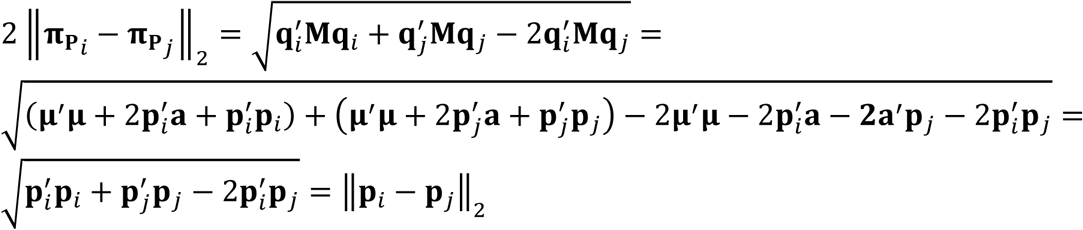

